# Is it chaotic? Chicken Linkage Disequilibrium and quantitative trait loci

**DOI:** 10.1101/2025.09.28.679118

**Authors:** Ehud Lipkin, Janet E. Fulton, Jacqueline Smith, David W. Burt, Morris Soller

## Abstract

Linkage disequilibrium (LD) underpins genome-wide association studies (GWAS) to map Quantitative Trait Loci Regions (QTLRs). A common yet misled assumption is that the closest sites to a significant marker are the prime candidates as the causative mutation. However, LD blocks are typically fragmented and intermingled. We studied chicken LD of mapped QTLRs for Marek’s Disease in numerous chicken populations. We estimated background LD by random sampling of non-syntenic or syntenic marker pairs from different or same chromosomes. We defined LD blocks as a group of markers located on the same chromosome having significant moderate (0.15 ≥ r^2^ < 0.7) or high (r^2^ ≥ 0.7) LD, regardless of distance and the number of mixed markers with no LD. Next, we studied QTLR LD by all SNP markers within the QTLRs. We found very complex, nearly chaotic, LD patterns, with fragmented and interdigitated LD blocks. Exceptional LD was found between QTLRs 4-5. LD blocks and protein networks shared by the two QTLRs suggest possible functional relationships. Thus, potential causative candidates could be found beyond non-LD sites, occasionally at a very large distance from a significant marker, and even in another QTLR. Multiple effects might make LD intrinsically chaotic, thus limiting GWAS informativity and repeatability. Chaotic complexity challenges the practical use of LD across populations, and must be accounted for while interpreting genetic mapping studies.

## Introduction

Linkage disequilibrium (LD) is defined as a high correlation among alleles of different genomic sites [Aerts et al., 2007; Ardlie et al., 2002; Reich et al., 2001; Wall and Pritchard, 2003a]. Useful LD for mapping and breeding operations indicates non-random association of alleles at the different loci [Aerts et al., 2007]. Appreciable LD is commonly found between close loci, decreasing rapidly with distance [e.g., Aerts et al., 2007; García-Gámez et al., 2012]. LD provides the basis for Genome Wide Association Studies (GWAS) to map quantitative trait loci (QTL) [Wall and Pritchard, 2003a]. However, patterns of LD are often noisy and very complex [Lipkin et al, 2023; Lipkin et al, 2024; Price et al., 2008; Wall and Pritchard, 2003 a, b]. e.g., long range LD (LRLD), can stretch over millions of base pairs [Aerts et al., 2007; Corbin et al., 2010; García-Gámez et al., 2012; Koch et al., 2013; Park, 2019; Peters et al., 2021; Price et al., 2008; Skelly et al., 2016; Wall and Pritchard, 2003 a, b]. That is, sites may have practically no LD with adjacent sites, but high LD with sites that are far apart. Suggested reasons for this phenomenon, predicted by population genetic models, are heterogeneity of recombination rate, and population history, size and structure [Wall and Pritchard, 2003 a, b].

Typically, LD pattern is characterized by LD blocks [Wall and Pritchard, 2003 a, b]. Various definitions were proposed for these blocks. Gabriel et al. (2002) used |D’| to define haplotype blocks as sets of consecutive sites with less than 5% evidence of recombination [Gabriel et al., 2002]. i.e., a haplotype block is defined as a range of consecutive sites where at least 95% have high LD among them, while no more than 5% of the sites have low LD with all others. However, sites with significant LD are often separated by much more than 5% of the sites with which they have practically no LD [Aerts et al., 2007; Allabi et al., 2005; Lipkin et al., 2013; Lipkin et al., 2023; Lipkin et al., 2024; Pérez O’Brien et al., 2014; Price et al., 2008; Wall and Pritchard, 2003 a, b]. As a result, LD blocks may actually be made of distant, apparent discontinuous sites.

The question of what defines continuity thus arises, as continuity is not universal. Rather, it is but a matter of the specific sites used and their quality control filtering. Thus, a set of consecutive sites in one study, could be broken by introducing more sites in another study, and vice versa - fragmented sites of one study could be consecutive in another if mid sites were not included.

LD complexity increases the difficulty of GWAS mapping, in particularly, challenging location of the specific causative element. A significant association may be found between a causative locus and markers beyond non-LD markers, far removed from the actual cause. This might falsely place the putative causative locus at a site far away from its actual location [Skelly et al., 2016].

On the other hand, LD may point at interactions between distant genomic regions (e.g., a receptor and a ligand, or a gene and a regulator), as well as co-evolution of different genomic regions affected by the same selection (natural or artificial). Specifically in the context of the present study, LD between QTL regions (QTLRs) may present functional relationships among them.

Thus, the objectives of the present study were to characterize LD within and between previously mapped Marek’s Disease (MD) resistance QTLRs [Smith et al., 2020], in multiple commercially utilized elite chicken populations. Here, the definition of LD blocks was not restricted as previously to consecutive markers with 5% low LD [e.g., Gabriel et al., 2002], but looked beyond it. This enabled study of LD over a long distance, and illustrated the need to account for it while interpretating mapping results. Furthermore, this study highlighted possible functional relationships between two of the QTLRs.

## Results

### Remapping QTLRs from Galgal4 to GRCg6a

Smith et al. (2020) mapped QTLRs based on the Galgal4 genomic build. Lipkin et al. (2024) adjusted the marker and QTLR locations to those provided by GRCg6a. The same 38 QTLRs found on Galgal4 were found with GRCg6a, with only negligible changes (Supplemental_Table_S1). The new markers and QTLR coordinates on GRCg6a were used for all analyses in the present study.

### LD significance in F_6_ based on non-syntenic and syntenic 600K SNP array

#### Background non-syntenic LD between random markers from different chromosomes

A sample of random non-syntenic marker pairs was used to assess the LD background in the F_6_ families. Non-syntenic r^2^ averaged 0.011 ± 0.016 (Table 1). Average family non-syntenic LDs had significant high negative correlation with the sample size (the number of individuals used to calculate a marker pair LD; i.e., Ind/pair in Table 1), r = −0.895 (p = 0.040). Combining all F_6_ families together, only 3.5% of the non-syntenic r^2^ values were above 0.05 (Supplemental_Table_S2). A single high LD of r^2^ = 0.991 was found in Family 2. Without any replication, this outlier was treated as a result of sampling variation. With a mean non-syntenic LD of 0.011 ± 0.016 (Table 1), any r^2^ > 0.043 is above the background LD and could be taken as significant. However, to be on the safe side, a conservative critical LD value of r^2^ ≥ 0.15 was chosen to define significant LD.

**Table 1.** Non-syntenic or syntenic LD between random 600K array marker pairs from different or the same chromosomes in the F_6_ families. Pairs - number of marker pairs; Ind/pairs - sample size: average number of individuals used to calculate a markers pair LD; Avg r^2^ - average r^2^ weighted by the number of pairs; SD - standard deviation; Min - minimum; Max - maximum.

| Markers | Statistic | 1 | 2 | 3 | 4 | 5 | Avg | Min | Max |
| --- | --- | --- | --- | --- | --- | --- | --- | --- | --- |
| Non-Syntenic | Pairs | 190,232 | 182,366 | 184,399 | 187,406 | 178,780 | 184,636.6 | 178,780 | 190,232 |
|  | Ind/pair | 176.6 | 231.7 | 354.0 | 219.5 | 200.7 | 236.3 | 177 | 354 |
| | Avg $r^2$ | 0.014 | 0.012 | 0.008 | 0.010 | 0.012 | 0.011 | 0.008 | 0.014 |
| | $r^2$ SD | 0.019 | 0.016 | 0.011 | 0.014 | 0.016 | 0.016 | 0.011 | 0.019 |
| | Min $r^2$ | 0.000 | 0.000 | 0.000 | 0.000 | 0.000 | 0.000 | 0.000 | 0.000 |
| | Max $r^2$ | 0.368 | 0.991 | 0.372 | 0.246 | 0.243 | 0.443 | 0.243 | 0.991 |
| Syntenic | Pairs | 207,770 | 200,411 | 200,866 | 204,453 | 195,323 | 201,764.6 | 195,323 | 207,770 |
|  | Ind/pair | 176.6 | 231.7 | 354.0 | 219.5 | 200.7 | 236.3 | 177 | 354 |
| | Avg $r^2$ | 0.119 | 0.112 | 0.109 | 0.110 | 0.119 | 0.114 | 0.109 | 0.119 |
| | $r^2$ SD | 0.221 | 0.218 | 0.216 | 0.218 | 0.224 | 0.220 | 0.216 | 0.224 |
| | Min $r^2$ | 0.000 | 0.000 | 0.000 | 0.000 | 0.000 | 0.000 | 0.000 | 0.000 |
| | Max $r^2$ | 1.000 | 1.000 | 1.000 | 1.000 | 1.000 | 1.000 | 1.000 | 1.000 |

#### Syntenic LD between random markers on the same chromosome

A sample of random syntenic marker pairs was used to assess the level of LD on the same chromosome (Table 1). Family means of syntenic r^2^ were all in close range around 0.11, averaging 0.114, ten times the 0.011 mean obtained for the non-syntenic LD. The negative correlation between sample size and LD was seen again (−0.745), albeit this time only approaching significance (p = 0.150). In all families, about two-thirds of the marker pairs had r^2^ ≤ 0.05, dropping rapidly thereafter (Supplemental_Table_S3). Pooled over all families, the proportion of r^2^ ≥ 0.15 (set above as a threshold of significance by the non-syntenic LD), was around 0.2, while about 5% of all r^2^ values were above 0.7. Hence, the range of 0.15 ≤ r^2^ < 0.7 was set as low to moderate LD and r^2^ ≥ 0.7 as high LD, and these values were used to define moderate and high LD blocks.

### LD within and between QTLRs in F_6_

A detailed analysis of the LD within and between all QTLRs on the same chromosome was carried out with all 600K markers located within the QTLRs, in all F_6_ families. Six chromosomes harbored more than one QTLR (Supplemental_Table_S1), thus enabling examination of significant LD between 21 pairs of QTLRs. When all families were pooled together, 0.374% of all QTLR pairs had high LD between QTLRs (Supplemental_Table_S4). Family 5 was an outlier, with a much lower number and proportion of across-QTLR LD. Further inspection did not identify any source of this difference. Hence, we have no explanation other than sampling variation. In all F_6_ families, high LDs were found between most pairs of QTLRs (Supplemental_Table_S5). An exceptionally large number high LD pairs was repeatedly found in all families between QTLRs 4 and 5 on Chromosome 1 (bolded in Supplemental_Table_S5), with a total of 159,413 high LDs combined (Supplemental_Table_S5f). Summing up across all families, this is 142 times more than the next largest number of high LDs between QTLRs (1,123 between QTLRs 6-7; Tables S5f, S6). Thus, the LD between these two QTLRs was indeed unusual.

### QTLR LD Blocks in the F_6_ population

High LD blocks were defined as stretches of markers on the same chromosome with high LD among them, even if not continuous, even if intermingled with more than 5% other markers with no LD, and even if they were very distant. Large, fragmented, and interdigitated LD blocks were found in all five F_6_ families over all six chromosomes examined. An example of fragmented interdigitated blocks is presented in Table 2a. Three high LD blocks are seen, all fragmented and all interdigitated with one another: Blocks 1, 2, 3 include 4, 3 and 2 fragments respectively. The fact that they are indeed genuine blocks, is shown in Tables 2 b-d. If the markers of the other blocks were not included in the analysis (e.g., not preselected or filtered out), then three clear unambiguous blocks would have been identified. Note that, should the upstream marker number 150 be the causative mutation, adjacent markers only kilobases away but with no LD whatsoever (r^2^ = 0.05) would not be affected, while more than half a megabase downstream, Marker number 337 with a complete LD of r^2^ = 1.000, would become significant. Thus, the problem that long-range fragmented LD blocks impose on locating a causative mutation is obvious.

**Table 2.**
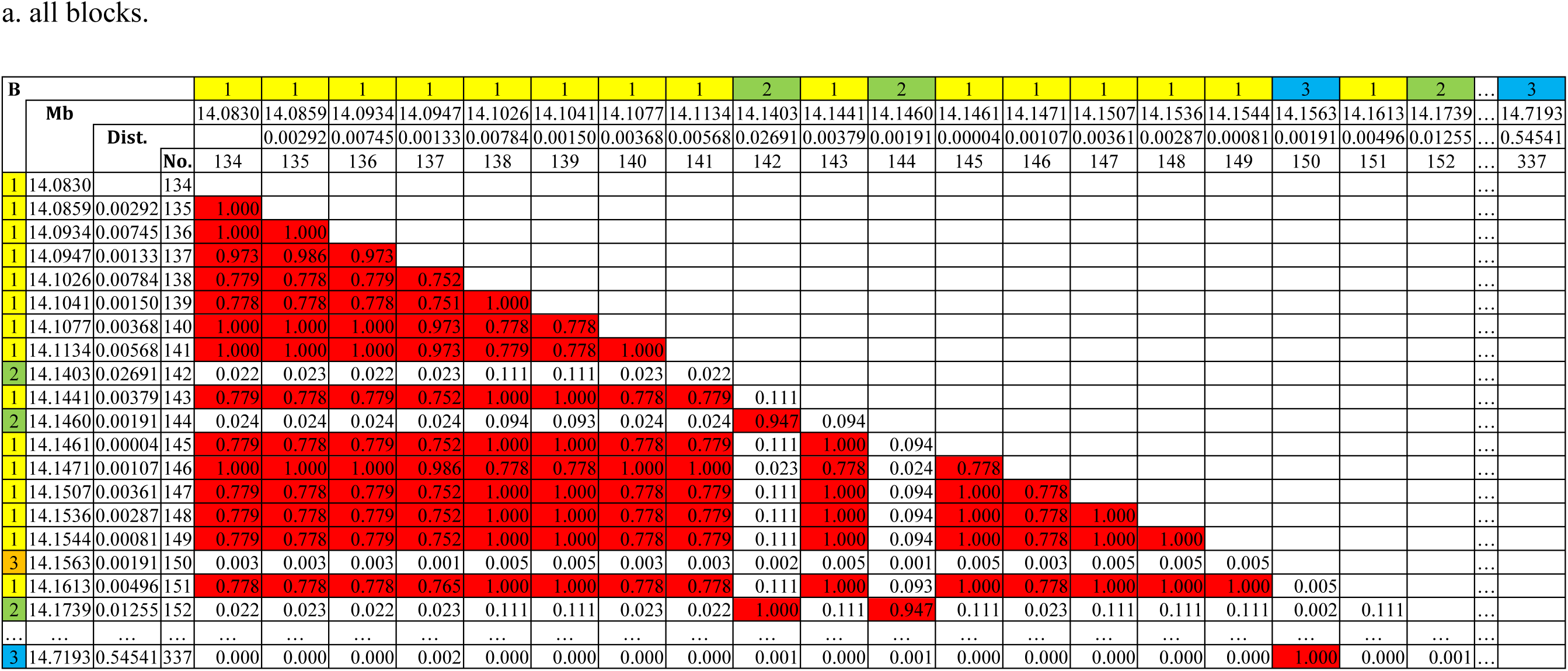
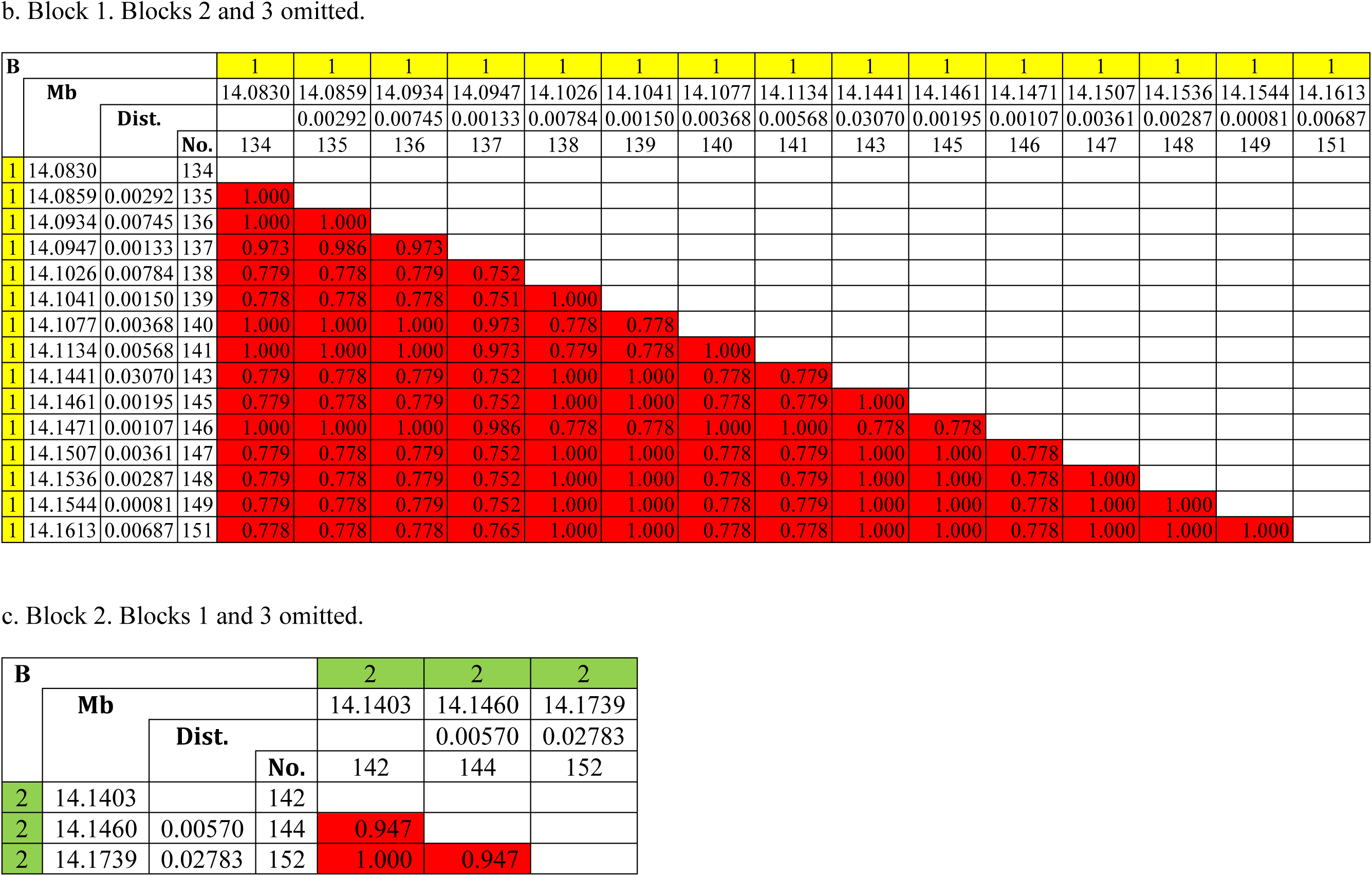
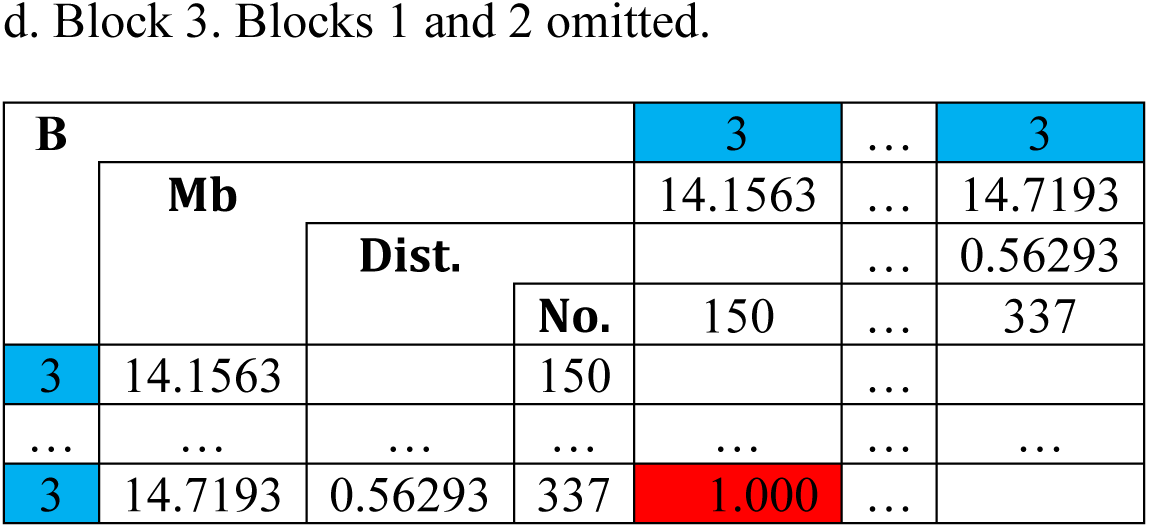
Fragmented interdigitated high LD blocks in QTLR 2 on chromosome 1 found in F_6_ Family 1. B, block serial number ordered by the location of the first marker of the block, coloured by blocks; Mb - marker location on GRCg6a in Mb; Dist. - distance between the marker and the previous one in Mb; … - large number of unpresented intermediate markers; No. - serial number of the marker; red - r^2^ ≥ 0.7; white r^2^ - r^2^ < 0.15. all blocks.

### LD Blocks extending QTLRs 4 and 5 in the F_6_ population

With all QTLR 600K markers, an exceptionally large number of long-range LD stretching from QTLR 4 to QTLR 5 was found in all five F_6_ families. As seen in Supplemental_Table_S7, for example, a high LD block composed of 412 markers extends from the first marker of QTLR 4, almost to the end of QTLR 5 (over 5.7Mb distance). Furthermore, moderate LD would extend the block all the way to the end of QTLR 5, encompassing over 7 Mb. Thus, the exceptional LD between QTLRs 4 and 5 indicated by the random marker sample, was confirmed by LD blocks based on all QTLR markers.

### LD within and between QTLRs in the eight lines

LD within and between all F_6_ QTLRs was also re-examined in eight Hy-Line elite egg production lines, using all QTLR lab markers [Smith et al., 2020]. Pooled over all lines, r^2^ values were obtained by markers from 22 QTLRs distributed over 15 chromosomes (Supplemental_Table_S8). Complex fragmented and interdigitated LD blocks were again found within and between QTLR elements (Supplemental_Fig_S11), similar to the above F_6_ blocks and to previous reports [Aerts et al., 2007; Allabi et al., 2005; Lipkin et al., 2013; Lipkin et al., 2023; Lipkin et al., 2024; Quinet et al., 2020]. Pooled over all lines, a total of 275 high LD blocks were found, stretching over 2 bp – 3.7 Mb (Tables S9, S10).

### LD blocks and QTLR elements in the eight lines

The QTLR lab markers used in the eight lines were selected to be within and around functional genomic elements [Smith et al., 2020]. This enabled examination of LD between QTLR elements. Complex LD blocks appear even between the elements within the same QTLR, and even within elements. Table 3, for example, presents three long noncoding RNAs (lncRNAs) in the same QTLR. Two blocks each consisting of 3 fragments are seen. The mid (green) lncRNA05 is split among the two blocks: some of its markers are in Block 1 with upstream lncRNA02, while others form Block 2 with the downstream lncRNA02. The two blocks are interdigitated within lncRNA05. Markers within each block had similar p-values, but were different from the other block. This match of p-values and LD blocks was found in all lines and QTLRs (Supplemental_Fig_S11 and further text below). More examples of similar complex LD blocks in protein coding genes within QTLRs are presented in the Supplemental materials.

**Table 3.**
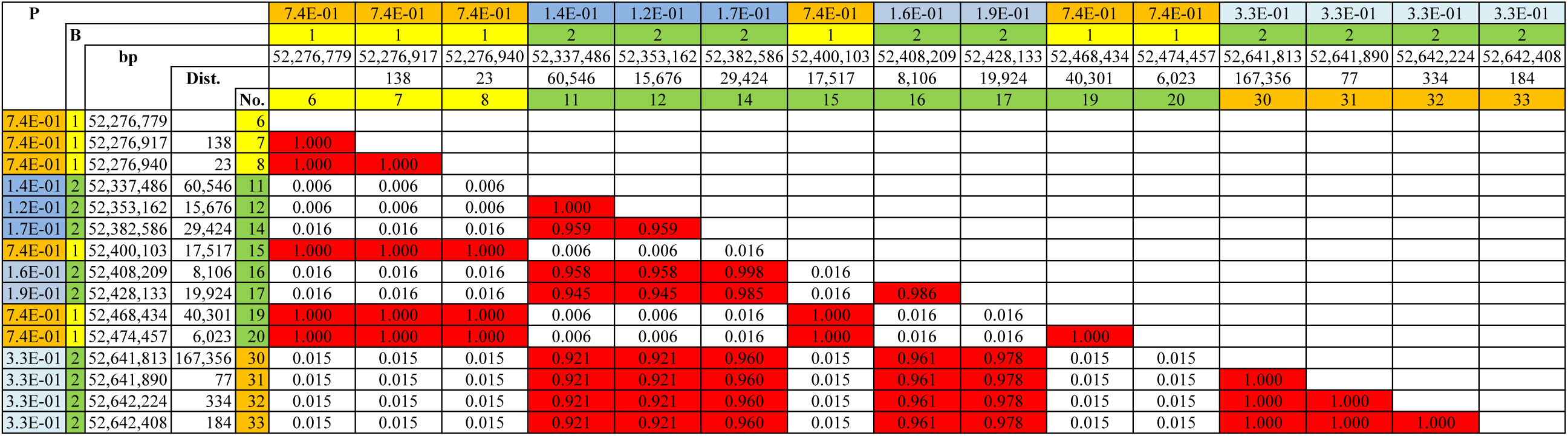
LD blocks across long noncoding RNAs (lncRNAs) in the same QTLR. Line WL3, QTLR 3 on chromosome 1. P - p-values obtained by the trend test [Smith et al., 2020], coloured by p-values; B - block serial number ordered by the location of the first marker, coloured by blocks; bp - marker location in GRCg6a; Dist. - distance between the present and previous marker in bp; No. - serial number the marker, coloured by the lncRNAs [Smith et al., 2020]: yellow, lncRNA02; green, lncRNA05; orange, lncRNA04; LD values: red r^2^ ≥ 0.7; white - r^2^ < 0.15.

### Is LD information applicable between populations?

Measuring QTLR LD in eight lines by the same markers enabled comparison of populations and assessment of the feasibility of using LD information from one population in another. Similar complexity of high LD blocks was found in all lines (Tables 4, S11). In light of the very complex LD pattern, comparing blocks among different populations is not a trivial task. This was done based on LD block boundaries [Schwartz et al., 2003], or individual marker pairs [Gabriel et al., 2002].

#### Blocks boundaries

Based merely on the block boundaries, a substantial overlap was found between high LD blocks (Supplemental_Table_S9, coloured in column ‘Distance’). However, once blocks within a population were allowed to overlap, the shared boundary statistic [Schwartz et al., 2003] would not be applicable. The fragmentation and the interdigitation (mixing) of blocks repeatedly break the overlap by non-overlapping mid markers and mixed LD blocks (Supplemental_Table_S11). That is, blocks from two lines may overlap at the start and the end, but not in the middle. See, for example, Table 4. Taking the boundaries, the two Block 1 of the two lines partially overlap. The situation, however, is more complicated than that. Careful examination shows that the overlap is broken by partially overlapping Block 2 in both lines, non-overlapping Block 3 in Line WL2 only, and non-overlapping markers with no LD to any other markers in Line WPR1. This almost chaotic complexity appears in all lines and blocks. Clearly the boundaries are not sufficient to define overlap of LD blocks.

**Table 4.** Comparing LD blocks among lines. Mixed high LD blocks in the start of QTLR 3 in lines WL2 and WPR1 (from Supplemental_Table_S11). Chr - chromosome; Location - location of the marker in bp in the GRCg6a chicken reference; Distance - base pairs between the marker and the previous one; Marker - coloured by the genomic element: Name - marker name including genomic element; no - serial number of the marker; Line blocks are numbered and coloured independently within a line (that is, Block 1 in one line is not necessarily Block 1 in the other lines; same colour in different lines does not imply same block); 0 - the marker had no high LD with any other marker; “-” - the marker was not informative in the line and thus was not tested in that line.

| Chr | Location | Distance | Marker |  | Line blocks |  |
| --- | --- | --- | --- | --- | --- | --- |
|  |  |  | Name | no. | WL2 | WPR1 |
| 1 | 52,276,776 |  | lncRNA02.01 | 1 | 1 | 1 |
| 1 | 52,276,779 | 3 | lncRNA02.02 | 2 | 2 | 2 |
| 1 | 52,276,917 | 138 | lncRNA02.03 | 3 | 2 | 2 |
| 1 | 52,276,940 | 23 | lncRNA02.04 | 4 | 3 | 0 |
| 1 | 52,276,945 | 5 | lncRNA02.05 | 5 | 1 | 1 |
| 1 | 52,277,046 | 101 | lncRNA02.06 | 6 | 1 | 1 |
| 1 | 52,337,486 | 60,440 | lncRNA05.20 | 7 | 2 | 3 |
| 1 | 52,353,162 | 15,676 | lncRNA05.21 | 8 | 3 | - |
| 1 | 52,370,944 | 17,782 | lncRNA05.22 | 9 | 1 | 4 |
| 1 | 52,382,586 | 11,642 | lncRNA05.23 | 10 | - | 0 |
| 1 | 52,400,103 | 17,517 | lncRNA05.24 | 11 | 0 | - |
| 1 | 52,408,209 | 8,106 | lncRNA05.25 | 12 | 1 | 3 |
| 1 | 52,428,133 | 19,924 | lncRNA05.26 | 13 | 0 | 3 |
| 1 | 52,453,441 | 25,308 | lncRNA05.27 | 14 | 4 | 0 |
| 1 | 52,468,434 | 14,993 | lncRNA05.28 | 15 | 3 | 4 |
| 1 | 52,474,457 | 6,023 | lncRNA05.29 | 16 | 3 | - |

#### Pairs of markers

A way to overcome the problems of comparing mixed blocks is to focus on the minimum possible block size, namely two markers only [Gabriel et al., 2002]. In the present study, two markers were assigned to the same high LD block if r^2^ ≥ 0.7, regardless of the distance or the presence of many mid markers with low LD. To this end, all marker pairs shared by two lines were counted (Supplemental_Table_S12, bottom left triangle), and pairs with high LD among them were identified (Supplemental_Table_S12, upper right triangle). Based on Supplemental_Table_S12, the proportions of the shared high LD pairs among all shared pairs were calculated (Table 5). For example, 127 marker pairs were shared by Lines WL1 and WL2, 40 of which had high LD (Supplemental_Table_S12), i.e. a proportion of 0.315 (Table 5). The proportion of shared high LD blocks consisting of two markers averaged 0.199 ± 0.131 (Table 5). This low sharing alone manifests the uniqueness of the LD pattern to each line, and consequently the difficulties in sharing LD information among the populations.

**Table 5.** Proportion of marker pairs with high LD, among all shared pairs (based on Supplemental_Table_S12). Avg, SD, Min and Max, average, standard deviation, minimum and maximum number of shared marker pairs between pairs of chicken lines; Total - all lines pooled together.

| Line | Marker pairs |  |  |  |  |  |  |  |  |  |  |  |
| --- | --- | --- | --- | --- | --- | --- | --- | --- | --- | --- | --- | --- |
|  | WL1 | WL2 | WL3 | WPR1 | WPR2 | WL4 | WL5 | RIR1 | Avg | SD | Min | Max |
| WL1 |  | 0.315 | 0.233 | 0.290 | 0.176 | 0.661 | 0.412 | 0.373 | 0.352 | 0.158 | 0.176 | 0.661 |
| WL2 |  |  | 0.136 | 0.081 | 0.069 | 0.221 | 0.245 | 0.115 | 0.169 | 0.093 | 0.069 | 0.315 |
| WL3 |  |  |  | 0.224 | 0.197 | 0.195 | 0.333 | 0.196 | 0.216 | 0.060 | 0.136 | 0.333 |
| WPR1 |  |  |  |  | 0.115 | 0.126 | 0.080 | 0.074 | 0.141 | 0.084 | 0.074 | 0.290 |
| WPR2 |  |  |  |  |  | 0.134 | 0.077 | 0.067 | 0.119 | 0.052 | 0.067 | 0.197 |
| WL4 |  |  |  |  |  |  | 0.130 | 0.155 | 0.232 | 0.193 | 0.126 | 0.661 |
| WL5 |  |  |  |  |  |  |  | 0.133 | 0.201 | 0.131 | 0.077 | 0.412 |
| RIR1 |  |  |  |  |  |  |  |  | 0.159 | 0.104 | 0.067 | 0.373 |
| **Total** |  |  |  |  |  |  |  |  | 0.199 | 0.131 | 0.067 | 0.661 |

### LD between QTLRs 4 and 5 in the eight elite lines

#### LD blocks

Informative markers located in both QTLRs 4-5 were found only in five of the eight lines (Supplemental_Table_S11). Furthermore, even in those lines there were only 2 - 4 informative markers per line in QTLR 4. Thus, there was limited information available for this analysis. Nevertheless, in accordance with the F_6_ analysis, moderate LD blocks were seen to extend over the two QTLRs. For example, two clear high LD blocks were found in Line WPR1, one in each QTLR (Table 6). The two high LD blocks had moderate LDs of r^2^ = 0.478 among them, thus forming one significant, albeit moderate, LD block. A similar pattern was found in three more lines (Supplemental_Fig_S11).

**Table 6.** LD in QTLRs 4-5. LD in Line WPR1 between lncRNA01 in QTLR 4 and the *CSTA* gene in QTLR 5. P - p-value obtained by e association trend test [Smith et al., 2020], coloured by p-values; Q - QTLR serial number (Supplemental_Table_S1), coloured by TLR; bp - marker location in GRCg6a; Dist. - distance between the present and previous marker in bp; No. - serial number of the arker, coloured by QTLR element [Smith et al., 2020]: yellow, lncRNA01; green, CSTA gene; LD values: Red - r^2^ ≥ 0.70; pink - 0.15 r^2^ < 0.70.

| P |  |  |  |  | 8.2E-02 | 8.2E-02 | 8.2E-02 | 8.2E-02 | 3.9E-01 | 3.9E-01 | 3.9E-01 |
| --- | --- | --- | --- | --- | --- | --- | --- | --- | --- | --- | --- |
|  | Q |  |  |  | 4 | 4 | 4 | 4 | 5 | 5 | 5 |
|  |  | bp |  |  | 72,307,283 | 72,307,369 | 72,307,616 | 72,307,630 | 77,442,145 | 77,444,900 | 77,444,931 |
|  |  |  | Dist. | No. |  | 86 | 247 | 14 | 5,134,515 | 2,755 | 31 |
|  |  |  |  |  | 34 | 36 | 37 | 38 | 42 | 44 | 45 |
| 8.2E-02 | 4 | 72,307,283 |  | 34 |  |  |  |  |  |  |  |
| 8.2E-02 | 4 | 72,307,369 | 86 | 36 | 1.000 |  |  |  |  |  |  |
| 8.2E-02 | 4 | 72,307,616 | 247 | 37 | 1.000 | 1.000 |  |  |  |  |  |
| 8.2E-02 | 4 | 72,307,630 | 14 | 38 | 1.000 | 1.000 | 1.000 |  |  |  |  |
| 3.9E-01 | 5 | 77,442,145 | 5,134,515 | 42 | 0.478 | 0.478 | 0.478 | 0.478 |  |  |  |
| 3.9E-01 | 5 | 77,444,900 | 2,755 | 44 | 0.478 | 0.478 | 0.478 | 0.478 | 1.000 |  |  |
| 3.9E-01 | 5 | 77,444,931 | 31 | 45 | 0.478 | 0.478 | 0.478 | 0.478 | 1.000 | 1.000 |  |

#### Bioinformatics analysis

Searching for a source of the exceptional LD between QTLRs 4 and 5, STRING network analysis found 10 and 68 genes in QTLRs 4 and 5, respectively (Supplemental_Table_S13). Seven networks of 2 to 19 genes were found (Figure 1 and Supplemental_Table_S13). Three of the networks were comprised of genes from both QTLRs (’Net’ 2, 3, 4 in Figure 1 and Supplemental_Table_S13). Except four of the genes in Network 4 in QTLR 5, all other 19 genes in those three cross-QTLRs networks were located in the F_6_ LD blocks extending over the two QTLRs (’+’ in the column ‘B4-5’ in Supplemental_Table_S13). Thus, genes from both QTLRs which had high LD among them, are also part of the same network.

**Figure 1.**
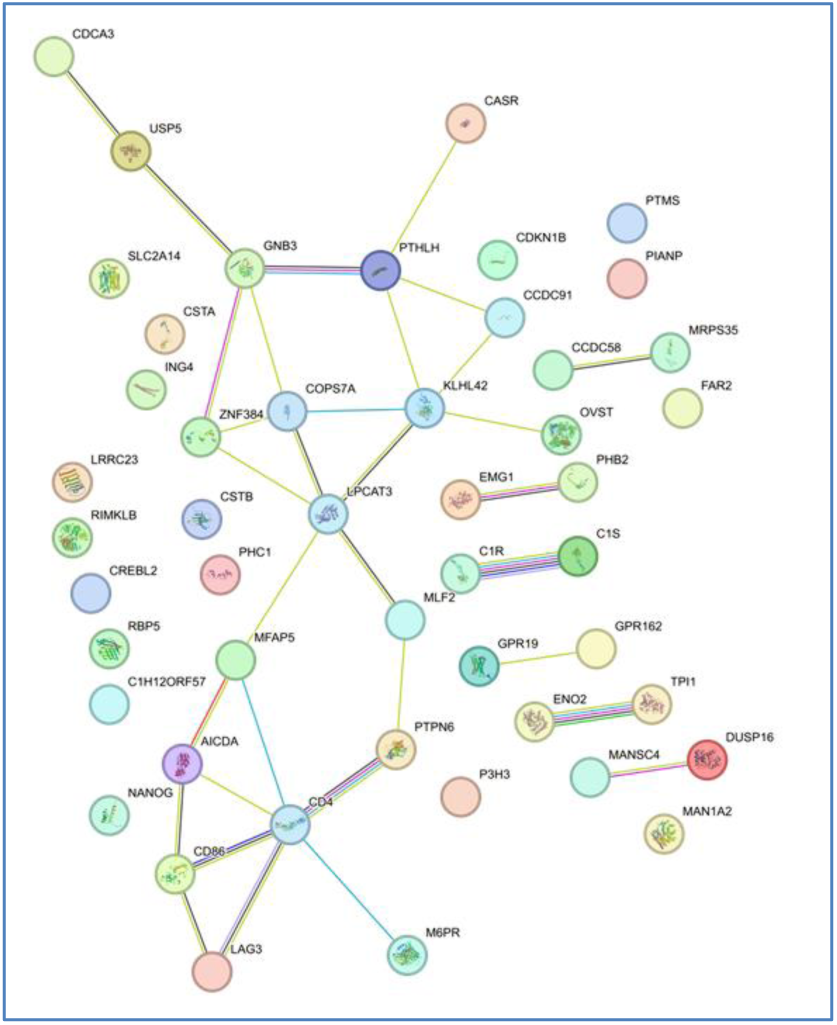
Genes from different QTLRs but in the same network. STRING network of genes under F_6_ QTLRs 4 and 5. Network nodes represent proteins; Coloured nodes, query proteins and first shell of interactors; Node content: empty, protein of unknown 3D structure; filled, some 3D structure is known or predicted; Known interactions: 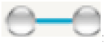; from curated databases; 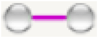, experimentally determined; Predicted interactions: 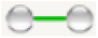, gene neighborhood; 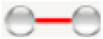, gene fusions; 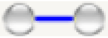, gene co-occurrence; Others: 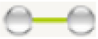, text mining; 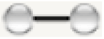, co-expression; 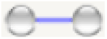, protein homology. The number of the QTLR is shown next to the gene, coloured by the QTLR (Supplemental_Table_S1).

## Discussion

Chicken LD was studied by available genotypes of an FSAIL F_6_ population and eight commercially relevant pure lines. The numerous populations and experimental designs allowed exploration of LD patterns and their repeatability.

### Background and significant LD

Background LD assessed by non-syntenic marker pairs was comparable to previous reports in chicken [Lipkin et al., 2013; Seo et al., 2018], but about 10 times higher than values reported for mammals; namely horse [Corbin et al., 2010], sheep [García-Gámez et al., 2012], and cattle [Khatkar et al., 2008; Sargolzaei et al., 2008]. These differences may be merely technical, representing differences in experimental design, sample size, sample variation, population history and/or structure. For instance, a population established by a cross will have lower LD than the one established by a few pure breed individuals; immigration or introgressions will present new alleles in a population, break relations between sites and reduce LD; intensity of selection will change LD. They could, however, represent true biological differences between birds and mammals, but the exact source of the difference is beyond the scope of the present study. The negative correlation between F_6_ family LD and the size of the population sample is in accord with a previous report in chicken [Pérez O’Brien et al., 2014], and the expectation of Sved (1971), thus reinforcing the experimental design and the results of the random marker samples.

### QTLR LD

A detailed LD analysis of all QTLRs on the same chromosome was carried out with all QTLR markers. Significant LD was found between most pairs of QTLRs in all five F_6_ families, with an exceptionally large amount of high LD between QTLRs 4 and 5 on chromosome 1 (below). Upon repeating in all F_6_ families, significant LD between QTLRs were found to be frequent, distributing in all chromosomes tested.

### LD blocks

Obviously, LD block analysis depends on the exact definition of the blocks. Here, LD blocks were defined as a group of markers located on the same chromosome having moderate or high LD with each other, regardless of the number of mid markers with low LD or the physical distance between the markers. This permissive definition is different from previous studies [e.g., Gabriel et al., 2002], and allowed us to look over the immediate horizon and identify complex mixed blocks beyond many non-LD markers over a long distance.

We show that the fragmented blocks are indeed genuine blocks, and if some of the markers in the fragmented blocks were not included in the analysis for technical reasons, then clear continuous blocks would have been identified. Needless to say, the opposite also holds: adding more markers may break previously identified continuous LD blocks.

The phenomenon of fragmented and interdigitated LD blocks was repeatedly found in all five F_6_ families and in the eight elite lines over a vast range of distances, from hundreds to megabases. The repeatability in multiple populations, and the agreement with previous studies by us and others [e.g., Aerts et al., 2007; Allabi et al., 2005; Lipkin et al., 2013; Lipkin et al., 2023; Lipkin et al., 2024; Pérez O’Brien et al., 2014; Wall and Pritchard, 2003 a, b], strengthens the notion that the extreme complex LD pattern of long range, fragmented and mixed blocks is a genuine and frequent biological phenomenon.

Interestingly, the F_6_ population was established from a cross of two lines and was designed to fragment the genome for high-resolution QTL mapping and thus reduce LD. The lines were pure, highly selected, and thus expected to have higher LD. Nevertheless, highly complex LD blocks were found in both datasets.

The complexity of high LD blocks found in all lines, challenges the comparison of blocks among different populations. Due to intermingled blocks, the boundaries of the LD blocks are not enough to define their overlap. Even reducing the blocks to the minimum possible size of two markers, still resulted in low sharing and large differences among the lines. Thus, using LD information from one population to another is very limited, if at all practical.

### LD between QTLRs 4 and 5

Repeating in all F_6_ families, an exceptionally large number of marker pairs with high LD were found between QTLRs 4 and 5. Accordingly, high LD blocks extending over both QTLRs were found. Similarly, moderate LD blocks between QTLRs 4 and 5 were also found in four of the pure lines. This exceptional LD is not due to the short distance between the two QTLRs: the pairs QTLRs 9-10 (distance of 2.17 Mb; Supplemental_Table_S1) and 24-23 (1.21 Mb) were closer than QTLRs 4-5 (2.24 Mb), with very little high LD (Tables S5, S6). The distance in other QTLR pairs was not much larger, with far smaller numbers of LDs among them (Tables S5, S6). Nor is it due to the location in the center of a chicken macrochromosome (chromosomes 1, 2, 4, 5 in this analysis). Recombination breaks LRLD, and the sub-telomeric regions of the macrochromosomes are known to exhibit very low recombination [Malinovskaya et al., 2019; Webster et al., 2006]. However, all other pairs of QTLRs are also located in sub-telomeric regions (Supplemental_Table_S1). For example, on the same region of chromosome 1, only a few high LD values were found between QTLRs 3-4 and 5-6, and no other pair of QTLRs on the three other macrochromosomes tested had a number of high LD values even remotely close to the number of high LDs found between QTLRs 4-5. Lastly, it is unlikely that such an exception is due to assembly errors [Qanbari et al., 2022; Utsunomiya et al., 2016]. Considering the very large number of cross-QTLR LDs, the number of potential errors would represent too numerous markers distributed over too large chromosomal regions.

### Bioinformatic analysis

We next wanted to understand what elements in QTLRs 4 and 5 could be causing such high LD over such large distances. Of course, being QTLRs for MD resistance, it is tempting to attribute this LD to co-selection for disease resistance. However, the question remains as to what makes this particular pair of QTLRs so different from all other pairs of QTLRs. STRING protein network analysis was used to reveal networks comprising genes from both QTLRs 4 and 5. Most of these genes were located within the cross QTLR high LD blocks. Thus, genes belonging to the same protein network and being located in the two different QTLRs, had significant LD in spite of the very large distance between them. These findings strengthen the possibility that it is functional relationships that cause the exceptional LD between QTLRs 4 and 5. These accords well with previous instances of finding functionally related genes on the same chromosomal region. Examples include the Major Histocompatibility Complex (MHC) on chicken chromosome 16, and the Regulators of Complement Activation cluster (RCA) on chromosome 26 [Allabi et al., 2005; Hosomichi et al., 2008; Oshiumi et al., 2005]. However, assessing the uniqueness of the LD between these particular QTLRs necessitates further study on the distribution of cross-QTLR networks and interactions among other pairs of QTLRs with less LD between them. Furthermore, assessing the real effect of the gene networks and interactions on the LD needs more molecular, quantitative and population studies, all of which are beyond the scope of the present study.

### Sources of LD complexity

The repeatability in different analyses, different datasets and numerous populations, and the agreement with previous reports, all strengthen the case for the present results. LD is complex, and indeed, almost chaotic. In general, the complexity of LD could stem from technical reasons such as sampling variation, mapping errors and genotyping errors. As presented above, marker preselection can merge or break LD blocks; population history, selection and structure all affect LD. In addition, LD could represent genuine biological relevance. Co-evolution of genomic sites as a result of natural or artificial selection may create long range LD between sites acting together; gene conversion could match alleles from distant sites; copy number variation may increase or decrease LD; founder effect and bottlenecks are expected to increase LD, non-random mating, epistasis, and immigration all may affect LD. Facing the extreme complexity of LD patterns, one cannot escape noting the chaotic nature of LD. This is possibly a result of the multiple effects shaping it, as multiple different effects are known to often result in chaotic behavior. In fact, chaotic complexity may as well be built-in LD behavior. If true, this feature would have a profound effect on GWAS and similar LD based operations.

### Implications for GWAS

Confirming past observations, the present results emphasize the need for caution while interpreting GWAS results. The basic assumption of GWAS is that a marker is significant possibly because it is the causative mutation or, much more likely, that it is in high LD with the true cause. However, in spite of the decay of LD with distance, often the closest markers have no practical LD with the significant one, while more distant markers have high LD. In the present study, pairs of markers as close as a few bp of each other had no LD and different p-values, while markers as distant as 12.5 Mb showed high LD and similar p-values. Thus, the mutation that causes the significance of a marker may not be a very close mutation, but could be a high LD site megabases away. In light of the exceptional LD and shared gene networks between QTLRs 4-5, the causative mutation beyond one QTLR may actually be in another QTLR. Thus, when inferring GWAS results, the complexity of the LD pattern must be taken into account. Last but not least, if chaotic complexity is indeed built-in to LD, this may well be the source of the poor repeatability of GWAS and other LD-dependent operations. If so, that poor repeatability is itself built-in to GWAS and other studies.

### Conclusions

The observed LD and fragmented interdigitated LD blocks imply that in genetic studies the causative element is not necessarily the closest, or even close at all, to the significant marker. Thus, mapping results and search for causative elements must consider the LD complexity. LD information in one population may have limited value in another. Multiple effects might drive LD to intrinsically chaotic behaviour, limiting GWAS informativity and repeatability.

## Methods

### Populations

The nine populations used by Smith et al. (2020), were used in the present study. These included five F_6_ families from a Full Sib Advanced Intercross Line (FSAIL) used to map the MD QTLRs tested here, and eight elite pure lines to test the QTLRs. The pure lines include the breeds White Leghorn (WL), Rhode Island Red (RIR), and White Plymouth Rock (WPR). Two of the WL lines were used for the development of the F_6_ FSAIL families.

*A priori*, it is expected that LD in the familial F_6_ population will be higher than in pure lines, as each family was initiated with only two parents [Heifetz et al., 2007]. Then again, this F_6_ population was designed to fragment the genome for high-resolution QTL mapping [Heifetz et al., 2007; Heifetz et al., 2009]. Thus, if at all, rather than inflated, LD in this population could in fact be under-estimated.

### Genotypes

The same autosomal genotypes obtained by Smith et al. (2020) to map and test QTLRs were used in the present study. F_6_ genotypes were obtained from the 600K Affymetrix SNP chicken array [Kranis et al., 2013], and genotypes of the eight pure lines were obtained from SNP markers developed in our lab (henceforth, “lab markers”) [Smith et al., 2020]. The only difference was that in F_6_, instead of a minimum MAF (Minor Allele Frequency) ≥ 0.01 used for the association tests by Smith et al. (2020), a threshold of 0.10 was used here for the LD analysis in the F_6_, to avoid spurious high LD due to rare alleles [Skelly et al., 2016; Wall and Pritchard, 2003b]. In the eight pure lines, only negligible differences were found between critical MAF of 0.01 and 0.10. Hence, to test LD with exactly the same markers used to test the QTLRs, the same MAF ≥ 0.01 as used by Smith et al. (2020), was used here too.

### Genome assemblies and remapping QTLRs

Smith et al. (2020) mapped QTLRs on the Galgal4 reference genome (Acc. No.: GCA_000002315.3). Lipkin et al., (2024) used the [Lift Genome Annotations](http://genome.ucsc.edu/cgi-bin/hgLiftOver) tool [Haeussler et al., 2019] within the [UCSC](http://genome.ucsc.edu/index.html) Browser to remap markers and QTLRs to the GRCg6a assembly (GRCg6a; Acc. No.: GCA_000002315.5). The new coordinates were used in the present study.

### Linkage disequilibrium (LD)

#### LD Measure

LD r^2^ was obtained using JMP Genomics software (JMP Genomics, Version 9, SAS Institute Inc., Cary, NC, USA, 1989–2019).

### LD between random non-syntenic or syntenic markers from different or the same chromosome in the F_6_ population

Background or typical LD were estimated each by two random combined samples with return of non-syntenic or syntenic marker pairs from respectively different or within a chromosome in each of the F_6_ families.

### LD and QTLRs

All markers located within the F_6_ MD QTLRs were used to measure QTLR LD in the F_6_ population with 600K markers (a total of 43,299,259 marker pairs), and in the eight lines with our lab markers (a total of 25,366 marker pairs). LD was measured between all possible pairs of markers on the same chromosome, including pairs from different QTLRs.

### LD blocks

Based on the distribution of the LD r^2^ values (see Results), moderate or high LD blocks were defined as a group of markers located on the same chromosome having respective LD of 0.15 > r^2^ ≤ 0.7 or r^2^ ≥ 0.7. This definition was applied even if many markers with low LD were mixed among the LD markers, and even if a large distance separated them. This definition allowed identification of long, fragmented and interdigitated blocks.

### Bioinformatics analysis

To investigate the genes underlying the exceptional LD between QTLRs 4 and 5 (see Results), the BioMart tool within the Ensembl database (https://www.ensembl.org/info/data/biomart/index.html) was used to identify genes within the QTLRs. The identified genes were then subjected to network analysis using the STRING database (v11) [Jensen et al., 2009], which provides an overview of known protein interactions.

## Data access

Data generated in this study have been submitted to the European Nucleotide Archive (ENA) at EMBL-EBI under study accession numbers PRJEB39142 (WGS) and PRJEB39361 (RNAseq).

## Acknowledgements

This research was funded by the Biotechnology and Biological Sciences Research Council (BBSRC; grant number BB/K006916/1). For the purpose of open access, the author has applied a creative commons attribution (CC BY) licence to any author accepted manuscript version arising.

