## Supplementals for "Is it chaotic? Chicken Linkage Disequilibrium and quantitative trait loci"

### Look over the horizon - chicken linkage disequilibrium and quantitative trait loci

#### SUPPLEMENTAL MATERIALS

**Supplemental_Table_S1.** Remapping QTLRs from Smith et al. (2020) on the chicken GRCg6a reference (Supplemental_Table_S6 in Lipkin et al., 2024). Q - ordinal number of the QTLR [Smith et al., 2020]; C - chromosome; Start/End - location of the first and last SNP of the QTLR on the GRCg6a genome build; Length - size of the QTLR (bp); Distance, base pairs between the first bp of the QTLR and last bp of the previous QTLR on the same chromosome.

| Q | C | Start | End | Length | Distance | Q | C | Start | End | Length | Distance |
|---|---|---|---|---|---|---|---|---|---|---|---|
| 1 | 1 | 9,510,148 | 9,902,036 | 391,889 | | 20 | 5 | 8,388,371 | 8,967,466 | 579,096 | |
| 2 | 1 | 13,994,599 | 14,950,768 | 956,170 | 4,092,563 | 21 | 5 | 19,753,005 | 20,610,009 | 857,005 | 10,785,539 |
| 3 | 1 | 52,166,588 | 52,643,244 | 476,657 | 37,215,820 | 22 | 6 | 3,323,132 | 3,946,659 | 623,528 | |
| 4 | 1 | 71,892,917 | 73,277,481 | 1,384,565 | 19,249,673 | 23 | 6 | 30,954,349 | 31,233,344 | 278,996 | 27,007,690 |
| 5 | 1 | 75,513,671 | 79,029,197 | 3,515,527 | 2,236,190 | 24 | 6 | 32,440,880 | 32,888,648 | 447,769 | 1,207,536 |
| 6 | 1 | 93,533,567 | 93,853,587 | 320,021 | 14,504,370 | 25 | 7 | 13,563,779 | 16,986,311 | 3,422,533 | |
| 7 | 1 | 103,738,415 | 106,416,920 | 2,678,506 | 9,884,828 | 26 | 10 | 1,025,523 | 2,668,959 | 1,643,437 | |
| 8 | 1 | 111,372,866 | 112,400,685 | 1,027,820 | 4,955,946 | 27 | 11 | 7,912,510 | 8,959,749 | 1,047,240 | |
| 9 | 1 | 171,680,812 | 174,306,953 | 2,626,142 | 59,280,127 | 28 | 12 | 9,414,714 | 9,845,036 | 430,323 | |
| 10 | 1 | 176,474,702 | 177,748,402 | 1,273,701 | 2,167,749 | 29 | 13 | 11,756,937 | 13,566,822 | 1,809,886 | |
| 11 | 1 | 196,152,404 | 196,750,875 | 598,472 | 18,404,002 | 30 | 14 | 8,499,374 | 9,745,708 | 1,246,335 | |
| 12 | 2 | 48,621 | 959,053 | 910,433 | | 31 | 14 | 13,542,085 | 15,384,231 | 1,842,147 | 3,796,377 |
| 13 | 2 | 45,786,534 | 46,247,754 | 461,221 | 44,827,481 | 32 | 16 | 1,852,095 | 2,669,032 | 816,938 | |
| 14 | 2 | 105,791,822 | 109,334,178 | 3,542,357 | 59,544,068 | 33 | 17 | 3,808,082 | 5,932,858 | 2,124,777 | |
| 15 | 2 | 125,532,963 | 127,219,187 | 1,686,225 | 16,198,785 | 34 | 18 | 3,221,049 | 4,118,252 | 897,204 | |
| 16 | 2 | 139,198,404 | 140,160,087 | 961,684 | 11,979,217 | 35 | 24 | 4,160,414 | 5,498,172 | 1,337,759 | |
| 17 | 3 | 108,593,746 | 109,643,999 | 1,050,254 | | 36 | 26 | 4,438,584 | 5,002,302 | 563,719 | |
| 18 | 4 | 8,328,709 | 11,309,259 | 2,980,551 | | 37 | 27 | 3,930,559 | 4,689,821 | 759,263 | |
| 19 | 4 | 84,829,085 | 89,057,374 | 4,228,290 | 73,519,826 | 38 | 28 | 1,447,725 | 1,687,264 | 239,540 | |

**Supplemental_Table_S2.** Non-syntenic LD. Distribution of non-syntenic LD values among random markers from different chromosomes in the $F_6$ families. $r^2$ >, sum of frequencies of $r^2 > 0.05$, 0.10, 0.15, respectively.

| $r^2$ | Family | | | | | All |
|---|---|---|---|---|---|---|
| | 1 | 2 | 3 | 4 | 5 | |
| ≤0.05 | 178,786 | 175,751 | 182,181 | 182,353 | 172,057 | 891,128 |
| >0.05 - ≤0.10 | 10,225 | 6,085 | 2,165 | 4,820 | 6,317 | 29,612 |
| >0.10 - ≤0.15 | 1,072 | 463 | 48 | 223 | 378 | 2,184 |
| >0.15 - ≤0.20 | 127 | 44 | 4 | 6 | 26 | 207 |
| >0.20 - ≤0.25 | 20 | 13 | 0 | 4 | 2 | 39 |
| >0.25 - ≤0.30 | 1 | 2 | 0 | 0 | 0 | 3 |
| >0.30 - ≤0.35 | 0 | 2 | 0 | 0 | 0 | 2 |
| >0.35 - ≤0.40 | 1 | 2 | 1 | 0 | 0 | 4 |
| >0.40 - ≤0.45 | 0 | 1 | 0 | 0 | 0 | 1 |
| >0.45 - ≤0.50 | 0 | 2 | 0 | 0 | 0 | 2 |
| >0.50 - ≤0.55 | 0 | 0 | 0 | 0 | 0 | 0 |
| >0.55 - ≤0.60 | 0 | 0 | 0 | 0 | 0 | 0 |
| >0.60 - ≤0.65 | 0 | 0 | 0 | 0 | 0 | 0 |
| >0.65 - ≤0.70 | 0 | 0 | 0 | 0 | 0 | 0 |
| >0.70 - ≤0.75 | 0 | 0 | 0 | 0 | 0 | 0 |
| >0.75 - ≤0.80 | 0 | 0 | 0 | 0 | 0 | 0 |
| >0.80 - ≤0.85 | 0 | 0 | 0 | 0 | 0 | 0 |
| >0.85 - ≤0.90 | 0 | 0 | 0 | 0 | 0 | 0 |
| >0.90 - ≤0.95 | 0 | 0 | 0 | 0 | 0 | 0 |
| >0.95 - ≤1.00 | 0 | 1 | 0 | 0 | 0 | 1 |
| Sum | 190,232 | 182,366 | 184,399 | 187,406 | 178,780 | 923,183 |
| $r^2 > 0.05$ | 0.06017 | 0.03627 | 0.01203 | 0.02696 | 0.03760 | 0.03472 |
| $r^2 > 0.10$ | 0.00642 | 0.00291 | 0.00029 | 0.00124 | 0.00227 | 0.00265 |
| $r^2 > 0.15$ | 0.00078 | 0.00037 | 0.00003 | 0.00005 | 0.00016 | 0.00028 |

**Supplemental_Table_S3.** Distribution of syntenic random LD values among the $F_6$ families. All - all families combined; $r^2$ >, sum of frequencies of $r^2 > 0.15$, 0.70, respectively.

| $r^2$ | Family | | | | | All |
|---|---|---|---|---|---|---|
| | 1 | 2 | 3 | 4 | 5 | |
| ≤ 0.05 | 0.6429 | 0.6778 | 0.6920 | 0.6920 | 0.6920 | 0.6707 |
| >0.05 - ≤0.10 | 0.1036 | 0.0898 | 0.0757 | 0.0757 | 0.0757 | 0.0879 |
| >0.10 - ≤0.15 | 0.0495 | 0.0451 | 0.0423 | 0.0423 | 0.0423 | 0.0457 |
| >0.15 - ≤0.20 | 0.0351 | 0.0277 | 0.0292 | 0.0292 | 0.0292 | 0.0315 |
| >0.20 - ≤0.25 | 0.0242 | 0.0192 | 0.0221 | 0.0221 | 0.0221 | 0.0219 |
| >0.25 - ≤0.30 | 0.0180 | 0.0165 | 0.0171 | 0.0171 | 0.0171 | 0.0174 |
| >0.30 - ≤0.35 | 0.0165 | 0.0138 | 0.0152 | 0.0152 | 0.0152 | 0.0152 |
| >0.35 - ≤0.40 | 0.0120 | 0.0110 | 0.0120 | 0.0120 | 0.0120 | 0.0122 |
| >0.40 - ≤0.45 | 0.0101 | 0.0101 | 0.0103 | 0.0103 | 0.0103 | 0.0102 |
| >0.45 - ≤0.50 | 0.0094 | 0.0092 | 0.0081 | 0.0081 | 0.0081 | 0.0091 |
| >0.50 - ≤0.55 | 0.0076 | 0.0087 | 0.0079 | 0.0079 | 0.0079 | 0.0080 |
| >0.55 - ≤0.60 | 0.0064 | 0.0079 | 0.0064 | 0.0064 | 0.0064 | 0.0070 |
| >0.60 - ≤0.65 | 0.0066 | 0.0076 | 0.0072 | 0.0072 | 0.0072 | 0.0069 |
| >0.65 - ≤0.70 | 0.0067 | 0.0069 | 0.0062 | 0.0062 | 0.0062 | 0.0066 |
| >0.70 - ≤0.75 | 0.0056 | 0.0070 | 0.0066 | 0.0066 | 0.0066 | 0.0062 |
| >0.75 - ≤0.80 | 0.0061 | 0.0062 | 0.0063 | 0.0063 | 0.0063 | 0.0060 |
| >0.80 - ≤0.85 | 0.0060 | 0.0063 | 0.0067 | 0.0067 | 0.0067 | 0.0063 |
| >0.85 - ≤0.90 | 0.0074 | 0.0068 | 0.0065 | 0.0065 | 0.0065 | 0.0068 |
| >0.90 - ≤0.95 | 0.0080 | 0.0073 | 0.0082 | 0.0082 | 0.0082 | 0.0081 |
| >0.95 - ≤1.00 | 0.0180 | 0.0151 | 0.0139 | 0.0139 | 0.0139 | 0.0162 |
| Sum | 1.0000 | 1.0000 | 1.0000 | 1.0000 | 1.0000 | 1.0000 |
| $r^2 > 0.15$ | 0.2040 | 0.1872 | 0.1900 | 0.1886 | 0.2089 | 0.1957 |
| $r^2 > 0.70$ | 0.0512 | 0.0486 | 0.0482 | 0.0486 | 0.0518 | 0.0497 |

**Supplemental_Table_S4.** QTLR LD. LD of all 600K marker pairs in and between QTLRs in the $F_6$ families. Number of pairs tested and sum of QTLR high LDs of $r^2 \geq 0.7$ found by SNP arrays in the $F_6$ families. All possible $F_6$ array pairs in and between QTLRs on the six chromosomes with more than one QTLR. Chr - chromosome; All - all families combined; Across QTLRs - high LD ($r^2 \geq 0.7$) of marker pairs starting upstream or within the upstream QTLR, ending within or downstream the downstream QTLR: N - number of pairs; % - proportion among all pairs.

| **Pairs** | **Chr** | **Family** | | | | | **All** |
|---|---|---|---|---|---|---|---|
| | | **1** | **2** | **3** | **4** | **5** | |
| **All** | 1 | 5,887,596 | 6,898,755 | 5,380,840 | 5,700,376 | 5,822,578 | 29,690,145 |
| | 2 | 1,046,181 | 1,113,778 | 952,890 | 1,017,451 | 689,725 | 4,820,025 |
| | 4 | 1,007,490 | 1,059,240 | 994,755 | 940,506 | 1,008,910 | 5,010,901 |
| | 5 | 108,345 | 110,685 | 42,195 | 103,285 | 108,345 | 472,855 |
| | 6 | 141,246 | 174,936 | 112,101 | 129,286 | 139,128 | 696,697 |
| | 14 | 440,391 | 675,541 | 424,581 | 533,028 | 535,095 | 2,608,636 |
| | Sum | 8,631,249 | 10,032,935 | 7,907,362 | 8,423,932 | 8,303,781 | 43,299,259 |
| **Across QTLRs** | N | 39,344 | 54,866 | 33,278 | 33,997 | 347 | 161,832 |
| | % | 0.456% | 0.547% | 0.421% | 0.404% | 0.004% | 0.374% |

**Supplemental_Table_S5. High LD values of random pairs of markers between QTLRs in the five $F_6$ families.** Number of 600K $F_6$ high LDs of $r^2 \geq 0.7$, starting upstream or within the upstream QTLR and ending within or downstream the downstream QTLR. C - chromosome separated by thick lines; Q - serial number of the $F_6$ MD QTLR (Supplemental_Table_S1); Mb - distance between the QTLR and the previous QTLR on the same chromosome; bolded - number of LDs between QTLRs 4-5.

a. Family 1.

| C | | | 1 | 1 | 1 | 1 | 1 | 1 | 1 | 1 | 1 | 1 | 1 | 2 | 2 | 2 | 2 | 2 | 4 | 4 | 5 | 5 | 6 | 6 | 6 | 14 | 14 |
|---|---|---|---|---|---|---|---|---|---|---|---|---|---|---|---|---|---|---|---|---|---|---|---|---|---|---|---|
| | Q | | 1 | 2 | 3 | 4 | 5 | 6 | 7 | 8 | 9 | 10 | 11 | 12 | 13 | 14 | 15 | 16 | 18 | 19 | 20 | 21 | 22 | 23 | 24 | 30 | 31 |
| | | Mb | | 4.1 | 37.2 | 19.2 | 2.2 | 14.5 | 9.9 | 5.0 | 59.3 | 2.2 | 18.4 | | 44.8 | 59.5 | 16.2 | 12.0 | | 73.5 | | 10.8 | | 27.0 | 1.2 | | 3.8 |
| 1 | 1 | | | | | | | | | | | | | | | | | | | | | | | | | | |
| 1 | 2 | 4.1 | 0 | | | | | | | | | | | | | | | | | | | | | | | | |
| 1 | 3 | 37.2 | 1 | 0 | | | | | | | | | | | | | | | | | | | | | | | |
| 1 | 4 | 19.2 | 1 | 3 | 1 | | | | | | | | | | | | | | | | | | | | | | |
| 1 | 5 | 2.2 | 0 | 4 | 3 | 39,114 | | | | | | | | | | | | | | | | | | | | | |
| 1 | 6 | 14.5 | 0 | 0 | 0 | 1 | 1 | | | | | | | | | | | | | | | | | | | | |
| 1 | 7 | 9.9 | 1 | 1 | 4 | 6 | 11 | 1 | | | | | | | | | | | | | | | | | | | |
| 1 | 8 | 5.0 | 0 | 1 | 0 | 1 | 5 | 0 | 2 | | | | | | | | | | | | | | | | | | |
| 1 | 9 | 59.3 | 0 | 3 | 2 | 6 | 16 | 0 | 10 | 4 | | | | | | | | | | | | | | | | | |
| 1 | 10 | 2.2 | 0 | 2 | 2 | 4 | 9 | 0 | 10 | 3 | 8 | | | | | | | | | | | | | | | | |
| 1 | 11 | 18.4 | 0 | 0 | 1 | 0 | 0 | 0 | 0 | 0 | 1 | 1 | | | | | | | | | | | | | | | |
| 2 | 12 | | | | | | | | | | | | | | | | | | | | | | | | | | |
| 2 | 13 | 44.8 | | | | | | | | | | | | 0 | | | | | | | | | | | | | |
| 2 | 14 | 59.5 | | | | | | | | | | | | 13 | 0 | | | | | | | | | | | | |
| 2 | 15 | 16.2 | | | | | | | | | | | | 8 | 0 | 11 | | | | | | | | | | | |
| 2 | 16 | 12.0 | | | | | | | | | | | | 3 | 0 | 13 | 4 | | | | | | | | | | |
| 4 | 18 | | | | | | | | | | | | | | | | | | | | | | | | | | |
| 4 | 19 | 73.5 | | | | | | | | | | | | | | | | | 26 | | | | | | | | |
| 5 | 20 | | | | | | | | | | | | | | | | | | | | | | | | | | |
| 5 | 21 | 10.8 | | | | | | | | | | | | | | | | | | | 1 | | | | | | |
| 6 | 22 | | | | | | | | | | | | | | | | | | | | | | | | | | |
| 6 | 23 | 27.0 | | | | | | | | | | | | | | | | | | | | | 0 | | | | |
| 6 | 24 | 1.2 | | | | | | | | | | | | | | | | | | | | | 1 | 1 | | | |
| 14 | 30 | | | | | | | | | | | | | | | | | | | | | | | | | | |
| 14 | 31 | 3.8 | | | | | | | | | | | | | | | | | | | | | | | | 19 | |

b. Family 2.

| **C** | | | **1** | **1** | **1** | **1** | **1** | **1** | **1** | **1** | **1** | **1** | **1** | **2** | **2** | **2** | **2** | **2** | **4** | **4** | **5** | **5** | **6** | **6** | **6** | **14** | **14** |
|---|---|---|---|---|---|---|---|---|---|---|---|---|---|---|---|---|---|---|---|---|---|---|---|---|---|---|---|
| | **Q** | | **1** | **2** | **3** | **4** | **5** | **6** | **7** | **8** | **9** | **10** | **11** | **12** | **13** | **14** | **15** | **16** | **18** | **19** | **20** | **21** | **22** | **23** | **24** | **30** | **31** |
| | | **Mb** | | **4.1** | **37.2** | **19.2** | **2.2** | **14.5** | **9.9** | **5.0** | **59.3** | **2.2** | **18.4** | | **44.8** | **59.5** | **16.2** | **12.0** | | **73.5** | | **10.8** | | **27.0** | **1.2** | | **3.8** |
| **1** | **1** | | | | | | | | | | | | | | | | | | | | | | | | | | |
| **1** | **2** | **4.1** | 0 | | | | | | | | | | | | | | | | | | | | | | | | |
| **1** | **3** | **37.2** | 0 | 0 | | | | | | | | | | | | | | | | | | | | | | | |
| **1** | **4** | **19.2** | 0 | 0 | 0 | | | | | | | | | | | | | | | | | | | | | | |
| **1** | **5** | **2.2** | 0 | 10 | 0 | **54,199** | | | | | | | | | | | | | | | | | | | | | |
| **1** | **6** | **14.5** | 0 | 4 | 0 | 0 | 9 | | | | | | | | | | | | | | | | | | | | |
| **1** | **7** | **9.9** | 0 | 17 | 0 | 3 | 56 | 16 | | | | | | | | | | | | | | | | | | | |
| **1** | **8** | **5.0** | 0 | 6 | 0 | 2 | 19 | 5 | 32 | | | | | | | | | | | | | | | | | | |
| **1** | **9** | **59.3** | 0 | 12 | 0 | 3 | 37 | 11 | 61 | 24 | | | | | | | | | | | | | | | | | |
| **1** | **10** | **2.2** | 0 | 4 | 0 | 3 | 22 | 3 | 32 | 13 | 21 | | | | | | | | | | | | | | | | |
| **1** | **11** | **18.4** | 0 | 4 | 1 | 0 | 8 | 2 | 15 | 5 | 12 | 3 | | | | | | | | | | | | | | | |
| **2** | **12** | | | | | | | | | | | | | | | | | | | | | | | | | | |
| **2** | **13** | **44.8** | | | | | | | | | | | | 0 | | | | | | | | | | | | | |
| **2** | **14** | **59.5** | | | | | | | | | | | | 20 | 0 | | | | | | | | | | | | |
| **2** | **15** | **16.2** | | | | | | | | | | | | 18 | 0 | 50 | | | | | | | | | | | |
| **2** | **16** | **12.0** | | | | | | | | | | | | 5 | 0 | 18 | 11 | | | | | | | | | | |
| **4** | **18** | | | | | | | | | | | | | | | | | | | | | | | | | | |
| **4** | **19** | **73.5** | | | | | | | | | | | | | | | | | 35 | | | | | | | | |
| **5** | **20** | | | | | | | | | | | | | | | | | | | | | | | | | | |
| **5** | **21** | **10.8** | | | | | | | | | | | | | | | | | | | 1 | | | | | | |
| **6** | **22** | | | | | | | | | | | | | | | | | | | | | | | | | | |
| **6** | **23** | **27.0** | | | | | | | | | | | | | | | | | | | | | 0 | | | | |
| **6** | **24** | **1.2** | | | | | | | | | | | | | | | | | | | | | 1 | 1 | | | |
| **14** | **30** | | | | | | | | | | | | | | | | | | | | | | | | | | |
| **14** | **31** | **3.8** | | | | | | | | | | | | | | | | | | | | | | | | 32 | |

38 c. Family 3.

| C | | | 1 | 1 | 1 | 1 | 1 | 1 | 1 | 1 | 1 | 1 | 1 | 2 | 2 | 2 | 2 | 2 | 4 | 4 | 5 | 5 | 6 | 6 | 6 | 14 | 14 |
|---|---|---|---|---|---|---|---|---|---|---|---|---|---|---|---|---|---|---|---|---|---|---|---|---|---|---|---|
| | **Q** | | **1** | **2** | **3** | **4** | **5** | **6** | **7** | **8** | **9** | **10** | **11** | **12** | **13** | **14** | **15** | **16** | **18** | **19** | **20** | **21** | **22** | **23** | **24** | **30** | **31** |
| | | **Mb** | | **4.1** | **37.2** | **19.2** | **2.2** | **14.5** | **9.9** | **5.0** | **59.3** | **2.2** | **18.4** | | **44.8** | **59.5** | **16.2** | **12.0** | | **73.5** | | **10.8** | | **27.0** | **1.2** | | **3.8** |
| **1** | **1** | | | | | | | | | | | | | | | | | | | | | | | | | | |
| **1** | **2** | **4.1** | 0 | | | | | | | | | | | | | | | | | | | | | | | | |
| **1** | **3** | **37.2** | 0 | 0 | | | | | | | | | | | | | | | | | | | | | | | |
| **1** | **4** | **19.2** | 0 | 0 | 0 | | | | | | | | | | | | | | | | | | | | | | |
| **1** | **5** | **2.2** | 0 | 1 | 0 | **33,181** | | | | | | | | | | | | | | | | | | | | | |
| **1** | **6** | **14.5** | 0 | 0 | 0 | 0 | 0 | | | | | | | | | | | | | | | | | | | | |
| **1** | **7** | **9.9** | 0 | 0 | 0 | 4 | 11 | 0 | | | | | | | | | | | | | | | | | | | |
| **1** | **8** | **5.0** | 0 | 0 | 0 | 0 | 0 | 0 | 0 | | | | | | | | | | | | | | | | | | |
| **1** | **9** | **59.3** | 0 | 0 | 0 | 1 | 2 | 0 | 3 | 2 | | | | | | | | | | | | | | | | | |
| **1** | **10** | **2.2** | 0 | 0 | 0 | 5 | 11 | 0 | 15 | 0 | 5 | | | | | | | | | | | | | | | | |
| **1** | **11** | **18.4** | 0 | 0 | 0 | 0 | 0 | 0 | 0 | 0 | 1 | 1 | | | | | | | | | | | | | | | |
| **2** | **12** | | | | | | | | | | | | | | | | | | | | | | | | | | |
| **2** | **13** | **44.8** | | | | | | | | | | | | 0 | | | | | | | | | | | | | |
| **2** | **14** | **59.5** | | | | | | | | | | | | 0 | 0 | | | | | | | | | | | | |
| **2** | **15** | **16.2** | | | | | | | | | | | | 0 | 0 | 0 | | | | | | | | | | | |
| **2** | **16** | **12.0** | | | | | | | | | | | | 0 | 0 | 0 | 1 | | | | | | | | | | |
| **4** | **18** | | | | | | | | | | | | | | | | | | | | | | | | | | |
| **4** | **19** | **73.5** | | | | | | | | | | | | | | | | | 24 | | | | | | | | |
| **5** | **20** | | | | | | | | | | | | | | | | | | | | | | | | | | |
| **5** | **21** | **10.8** | | | | | | | | | | | | | | | | | | | 0 | | | | | | |
| **6** | **22** | | | | | | | | | | | | | | | | | | | | | | | | | | |
| **6** | **23** | **27.0** | | | | | | | | | | | | | | | | | | | | | 0 | | | | |
| **6** | **24** | **1.2** | | | | | | | | | | | | | | | | | | | | | 2 | 0 | | | |
| **14** | **30** | | | | | | | | | | | | | | | | | | | | | | | | | | |
| **14** | **31** | **3.8** | | | | | | | | | | | | | | | | | | | | | | | | 8 | |

39

d. Family 4.

| C | | | 1 | 1 | 1 | 1 | 1 | 1 | 1 | 1 | 1 | 1 | 1 | 2 | 2 | 2 | 2 | 2 | 4 | 4 | 5 | 5 | 6 | 6 | 6 | 14 | 14 |
|---|---|---|---|---|---|---|---|---|---|---|---|---|---|---|---|---|---|---|---|---|---|---|---|---|---|---|---|
| | Q | | 1 | 2 | 3 | 4 | 5 | 6 | 7 | 8 | 9 | 10 | 11 | 12 | 13 | 14 | 15 | 16 | 18 | 19 | 20 | 21 | 22 | 23 | 24 | 30 | 31 |
| | | Mb | | 4.1 | 37.2 | 19.2 | 2.2 | 14.5 | 9.9 | 5.0 | 59.3 | 2.2 | 18.4 | | 44.8 | 59.5 | 16.2 | 12.0 | | 73.5 | | 10.8 | | 27.0 | 1.2 | | 3.8 |
| 1 | 1 | | | | | | | | | | | | | | | | | | | | | | | | | | |
| 1 | 2 | 4.1 | 0 | | | | | | | | | | | | | | | | | | | | | | | | |
| 1 | 3 | 37.2 | 0 | 0 | | | | | | | | | | | | | | | | | | | | | | | |
| 1 | 4 | 19.2 | 1 | 5 | 0 | | | | | | | | | | | | | | | | | | | | | | |
| 1 | 5 | 2.2 | 1 | 6 | 0 | **32,599** | | | | | | | | | | | | | | | | | | | | | |
| 1 | 6 | 14.5 | 0 | 0 | 0 | 0 | 0 | | | | | | | | | | | | | | | | | | | | |
| 1 | 7 | 9.9 | 0 | 3 | 0 | 5 | 35 | 1,106 | | | | | | | | | | | | | | | | | | | |
| 1 | 8 | 5.0 | 1 | 1 | 0 | 1 | 6 | 0 | 7 | | | | | | | | | | | | | | | | | | |
| 1 | 9 | 59.3 | 3 | 7 | 0 | 11 | 25 | 0 | 24 | 6 | | | | | | | | | | | | | | | | | |
| 1 | 10 | 2.2 | 0 | 1 | 0 | 5 | 28 | 0 | 24 | 5 | 18 | | | | | | | | | | | | | | | | |
| 1 | 11 | 18.4 | 0 | 0 | 1 | 0 | 0 | 0 | 0 | 0 | 1 | 1 | | | | | | | | | | | | | | | |
| 2 | 12 | | | | | | | | | | | | | | | | | | | | | | | | | | |
| 2 | 13 | 44.8 | | | | | | | | | | | | 0 | | | | | | | | | | | | | |
| 2 | 14 | 59.5 | | | | | | | | | | | | 0 | 0 | | | | | | | | | | | | |
| 2 | 15 | 16.2 | | | | | | | | | | | | 0 | 0 | 0 | | | | | | | | | | | |
| 2 | 16 | 12.0 | | | | | | | | | | | | 0 | 0 | 0 | 1 | | | | | | | | | | |
| 4 | 18 | | | | | | | | | | | | | | | | | | | | | | | | | | |
| 4 | 19 | 73.5 | | | | | | | | | | | | | | | | | 33 | | | | | | | | |
| 5 | 20 | | | | | | | | | | | | | | | | | | | | | | | | | | |
| 5 | 21 | 10.8 | | | | | | | | | | | | | | | | | | | 0 | | | | | | |
| 6 | 22 | | | | | | | | | | | | | | | | | | | | | | | | | | |
| 6 | 23 | 27.0 | | | | | | | | | | | | | | | | | | | | | 0 | | | | |
| 6 | 24 | 1.2 | | | | | | | | | | | | | | | | | | | | | 2 | 0 | | | |
| 14 | 30 | | | | | | | | | | | | | | | | | | | | | | | | | | |
| 14 | 31 | 3.8 | | | | | | | | | | | | | | | | | | | | | | | | 24 | |

e. Family 5.

| C | | | 1 | 1 | 1 | 1 | 1 | 1 | 1 | 1 | 1 | 1 | 1 | 2 | 2 | 2 | 2 | 2 | 4 | 4 | 5 | 5 | 6 | 6 | 6 | 14 | 14 |
|---|---|---|---|---|---|---|---|---|---|---|---|---|---|---|---|---|---|---|---|---|---|---|---|---|---|---|---|
| | Q | | 1 | 2 | 3 | 4 | 5 | 6 | 7 | 8 | 9 | 10 | 11 | 12 | 13 | 14 | 15 | 16 | 18 | 19 | 20 | 21 | 22 | 23 | 24 | 30 | 31 |
| | | Mb | | 4.1 | 37.2 | 19.2 | 2.2 | 14.5 | 9.9 | 5.0 | 59.3 | 2.2 | 18.4 | | 44.8 | 59.5 | 16.2 | 12.0 | | 73.5 | | 10.8 | | 27.0 | 1.2 | | 3.8 |
| 1 | 1 | | | | | | | | | | | | | | | | | | | | | | | | | | |
| 1 | 2 | 4.1 | 0 | | | | | | | | | | | | | | | | | | | | | | | | |
| 1 | 3 | 37.2 | 0 | 0 | | | | | | | | | | | | | | | | | | | | | | | |
| 1 | 4 | 19.2 | 0 | 0 | 0 | | | | | | | | | | | | | | | | | | | | | | |
| 1 | 5 | 2.2 | 0 | 0 | 0 | **320** | | | | | | | | | | | | | | | | | | | | | |
| 1 | 6 | 14.5 | 0 | 0 | 0 | 0 | 0 | | | | | | | | | | | | | | | | | | | | |
| 1 | 7 | 9.9 | 0 | 0 | 0 | 0 | 0 | 0 | | | | | | | | | | | | | | | | | | | |
| 1 | 8 | 5.0 | 0 | 0 | 0 | 0 | 0 | 0 | 0 | | | | | | | | | | | | | | | | | | |
| 1 | 9 | 59.3 | 0 | 0 | 0 | 0 | 0 | 0 | 0 | 0 | | | | | | | | | | | | | | | | | |
| 1 | 10 | 2.2 | 0 | 0 | 0 | 0 | 0 | 0 | 0 | 0 | 1 | | | | | | | | | | | | | | | | |
| 1 | 11 | 18.4 | 0 | 0 | 0 | 0 | 0 | 0 | 0 | 0 | 0 | 0 | | | | | | | | | | | | | | | |
| 2 | 12 | | | | | | | | | | | | | | | | | | | | | | | | | | |
| 2 | 13 | 44.8 | | | | | | | | | | | | 0 | | | | | | | | | | | | | |
| 2 | 14 | 59.5 | | | | | | | | | | | | 0 | 0 | | | | | | | | | | | | |
| 2 | 15 | 16.2 | | | | | | | | | | | | 0 | 0 | 0 | | | | | | | | | | | |
| 2 | 16 | 12.0 | | | | | | | | | | | | 0 | 0 | 0 | 0 | | | | | | | | | | |
| 4 | 18 | | | | | | | | | | | | | | | | | | | | | | | | | | |
| 4 | 19 | 73.5 | | | | | | | | | | | | | | | | | 22 | | | | | | | | |
| 5 | 20 | | | | | | | | | | | | | | | | | | | | | | | | | | |
| 5 | 21 | 10.8 | | | | | | | | | | | | | | | | | | | 0 | | | | | | |
| 6 | 22 | | | | | | | | | | | | | | | | | | | | | | | | | | |
| 6 | 23 | 27.0 | | | | | | | | | | | | | | | | | | | | | 0 | | | | |
| 6 | 24 | 1.2 | | | | | | | | | | | | | | | | | | | | | 0 | 0 | | | |
| 14 | 30 | | | | | | | | | | | | | | | | | | | | | | | | | | |
| 14 | 31 | 3.8 | | | | | | | | | | | | | | | | | | | | | | | | 4 | |

f. All families combined.

| C | | | 1 | 1 | 1 | 1 | 1 | 1 | 1 | 1 | 1 | 1 | 1 | 2 | 2 | 2 | 2 | 2 | 4 | 4 | 5 | 5 | 6 | 6 | 6 | 14 | 14 |
|---|---|---|---|---|---|---|---|---|---|---|---|---|---|---|---|---|---|---|---|---|---|---|---|---|---|---|---|
| | Q | | 1 | 2 | 3 | 4 | 5 | 6 | 7 | 8 | 9 | 10 | 11 | 12 | 13 | 14 | 15 | 16 | 18 | 19 | 20 | 21 | 22 | 23 | 24 | 30 | 31 |
| | | Mb | | 4.1 | 37.2 | 19.2 | 2.2 | 14.5 | 9.9 | 5.0 | 59.3 | 2.2 | 18.4 | | 44.8 | 59.5 | 16.2 | 12.0 | | 73.5 | | 10.8 | | 27.0 | 1.2 | | 3.8 |
| **1** | **1** | | | | | | | | | | | | | | | | | | | | | | | | | | |
| **1** | **2** | **4.1** | 0 | | | | | | | | | | | | | | | | | | | | | | | | |
| **1** | **3** | **37.2** | 1 | 0 | | | | | | | | | | | | | | | | | | | | | | | |
| **1** | **4** | **19.2** | 2 | 8 | 1 | | | | | | | | | | | | | | | | | | | | | | |
| **1** | **5** | **2.2** | 1 | 21 | 3 | **159,413** | | | | | | | | | | | | | | | | | | | | | |
| **1** | **6** | **14.5** | 0 | 4 | 0 | 1 | 10 | | | | | | | | | | | | | | | | | | | | |
| **1** | **7** | **9.9** | 1 | 21 | 4 | 18 | 113 | 1,123 | | | | | | | | | | | | | | | | | | | |
| **1** | **8** | **5.0** | 1 | 8 | 0 | 4 | 30 | 5 | 41 | | | | | | | | | | | | | | | | | | |
| **1** | **9** | **59.3** | 3 | 22 | 2 | 21 | 80 | 11 | 98 | 36 | | | | | | | | | | | | | | | | | |
| **1** | **10** | **2.2** | 0 | 7 | 2 | 17 | 70 | 3 | 81 | 21 | 53 | | | | | | | | | | | | | | | | |
| **1** | **11** | **18.4** | 0 | 4 | 3 | 0 | 8 | 2 | 15 | 5 | 15 | 6 | | | | | | | | | | | | | | | |
| **2** | **12** | | | | | | | | | | | | | | | | | | | | | | | | | | |
| **2** | **13** | **44.8** | | | | | | | | | | | | 0 | | | | | | | | | | | | | |
| **2** | **14** | **59.5** | | | | | | | | | | | | 33 | 0 | | | | | | | | | | | | |
| **2** | **15** | **16.2** | | | | | | | | | | | | 26 | 0 | 61 | | | | | | | | | | | |
| **2** | **16** | **12.0** | | | | | | | | | | | | 8 | 0 | 31 | 17 | | | | | | | | | | |
| **4** | **18** | | | | | | | | | | | | | | | | | | | | | | | | | | |
| **4** | **19** | **73.5** | | | | | | | | | | | | | | | | | 140 | | | | | | | | |
| **5** | **20** | | | | | | | | | | | | | | | | | | | | | | | | | | |
| **5** | **21** | **10.8** | | | | | | | | | | | | | | | | | | | 2 | | | | | | |
| **6** | **22** | | | | | | | | | | | | | | | | | | | | | | | | | | |
| **6** | **23** | **27.0** | | | | | | | | | | | | | | | | | | | | | 0 | | | | |
| **6** | **24** | **1.2** | | | | | | | | | | | | | | | | | | | | | 6 | 2 | | | |
| **14** | **30** | | | | | | | | | | | | | | | | | | | | | | | | | | |
| **14** | **31** | **3.8** | | | | | | | | | | | | | | | | | | | | | | | | 87 | |

**Supplemental_Table_S6.** LD between adjacent QTLRs on the same chromosome in the $F_6$ families. Distance and number of 600K high LD values of random pairs of markers between adjacent QTLRs on the same chromosome in all $F_6$ families. QTLR - serial number of the MD QTLR (Supplemental_Table_S1); Chr - chromosome; Length - size of the QTLR (bp); Distance - base pairs between the first bp of the QTLR and last bp of the previous QTLR on the same chromosome; LDs - number of high LDs ($r^2 \geq 0.7$), starting upstream or within the upstream QTLR, ending within or downstream of the downstream QTLR.

| QTLR | Chr | Length | Distance | LDs |
|---|---|---|---|---|
| 1 | 1 | 391,889 | | |
| 2 | 1 | 956,170 | 4,092,563 | 0 |
| 3 | 1 | 476,657 | 37,215,820 | 0 |
| 4 | 1 | 1,384,565 | 19,249,673 | 1 |
| 5 | 1 | 3,515,527 | 2,236,190 | 159,413 |
| 6 | 1 | 320,021 | 14,504,370 | 10 |
| 7 | 1 | 2,678,506 | 9,884,828 | 1,123 |
| 8 | 1 | 1,027,820 | 4,955,946 | 41 |
| 9 | 1 | 2,626,142 | 59,280,127 | 36 |
| 10 | 1 | 1,273,701 | 2,167,749 | 53 |
| 11 | 1 | 598,472 | 18,404,002 | 6 |
| 12 | 2 | 910,433 | | |
| 13 | 2 | 461,221 | 44,827,481 | 0 |
| 14 | 2 | 3,542,357 | 59,544,068 | 0 |
| 15 | 2 | 1,686,225 | 16,198,785 | 61 |
| 16 | 2 | 961,684 | 11,979,217 | 17 |
| 18 | 4 | 2,980,551 | | |
| 19 | 4 | 4,228,290 | 73,519,826 | 140 |
| 20 | 5 | 579,096 | | |
| 21 | 5 | 857,005 | 10,785,539 | 2 |
| 22 | 6 | 623,528 | | |
| 23 | 6 | 278,996 | 27,007,690 | 0 |
| 24 | 6 | 447,769 | 1,207,536 | 2 |
| 30 | 14 | 1,246,335 | | |
| 31 | 14 | 1,842,147 | 3,796,377 | 87 |

**Supplemental_Table_S7.** LD blocks in QTLRs 4-5. LD blocks shared by QTLRs 4 and 5 in $F_6$ Family 2. Q - QTLR serial number (Supplemental_Table_S1), coloured by QTLR; bp location in the GRCg6a reference in bp; Dist. - distance between the marker and the previous one in bp; … - large number of unpresented intermediate markers; No. - serial number of the marker; red $r^2$ - $r^2 \geq 0.7$; pink $r^2$ - $0.15 \leq r^2 < 0.7$; white $r^2$ - $r^2 < 0.15$.

| **Q** | | | | 4 | 4 | 4 | 4 | … | 5 | 5 | 5 | … | 5 | 5 | 5 |
|---|---|---|---|---|---|---|---|---|---|---|---|---|---|---|---|
| | **bp** | | | 71,892,917 | 71,895,755 | 71,895,827 | 71,898,143 | … | 77,550,920 | 77,559,651 | 77,560,137 | … | 79,021,461 | 79,022,071 | 79,025,156 |
| | | **Dist.** | | | 2,838 | 72 | 2,316 | … | 5,652,777 | 8,731 | 486 | … | 1,461,324 | 610 | 3,085 |
| | | | **No.** | 582 | 583 | 584 | 585 | … | 1057 | 1058 | 1059 | … | 1318 | 1319 | 1320 |
| 4 | 71,892,917 | | 582 | | | | | … | | | | … | | | |
| 4 | 71,895,755 | 2,838 | 583 | 1.000 | | | | … | | | | … | | | |
| 4 | 71,895,827 | 72 | 584 | 1.000 | 1.000 | | | … | | | | … | | | |
| 4 | 71,898,143 | 2,316 | 585 | 1.000 | 1.000 | 1.000 | | … | | | | … | | | |
| … | … | … | … | … | … | … | … | … | … | … | … | … | … | … | … |
| 5 | 77,550,920 | 5,652,777 | 1057 | 0.745 | 0.745 | 0.745 | 0.744 | … | | | | … | | | |
| 5 | 77,559,651 | 8,731 | 1058 | 0.745 | 0.745 | 0.745 | 0.744 | … | 1.000 | | | … | | | |
| 5 | 77,560,137 | 486 | 1059 | 0.000 | 0.000 | 0.000 | 0.000 | … | 0.000 | 0.000 | | … | | | |
| … | … | … | … | … | … | … | … | … | … | … | … | … | … | … | … |
| 5 | 79,021,461 | 1,461,324 | 1318 | 0.491 | 0.491 | 0.491 | 0.489 | … | 0.677 | 0.677 | 0.000 | … | | | |
| 5 | 79,022,071 | 610 | 1319 | 0.488 | 0.488 | 0.488 | 0.486 | … | 0.675 | 0.675 | 0.000 | … | 1.000 | | |
| 5 | 79,025,156 | 3,085 | 1320 | 0.472 | 0.472 | 0.472 | 0.471 | … | 0.656 | 0.656 | 0.001 | … | 0.972 | 0.972 | |

**Supplemental_Table_S8.** QTLR LD in the eight pure lines. The number of QTLR marker pairs used to analyze QTLR LD in the eight elite commercial egg production lines, by chromosome. Chr - chromosomes; QTLRs - QTLR serial numbers (Supplemental_Table_S1); Sum column - total number of marker pairs, by chromosomes; Sum line - Total number of marker pairs, by lines.

| Chr | QTLRs | Line | | | | | | | | Sum |
|---|---|---|---|---|---|---|---|---|---|---|
| | | WL1 | WL2 | WL3 | WPR1 | WPR2 | WL4 | WL5 | RIR1 | |
| 1 | 1,3,4,5,7,9,10,11 | 300 | 2,211 | 1,431 | 4,095 | 3,403 | 1,711 | 55 | 4,560 | 17,766 |
| 2 | 12,13 | 66 | 231 | 231 | 406 | 406 | 3 | 3 | 378 | 1,724 |
| 4 | 19 | 6 | 36 | 21 | 10 | 10 | 28 | 0 | 3 | 114 |
| 5 | 20,21 | 15 | 210 | 55 | 300 | 276 | 231 | 91 | 231 | 1,409 |
| 6 | 24 | 78 | 91 | 136 | 36 | 45 | 120 | 10 | 55 | 571 |
| 7 | 25 | 1 | 0 | 1 | 3 | 3 | 0 | 6 | 1 | 15 |
| 10 | 26 | 0 | 1 | 0 | 1 | 1 | 1 | 0 | 1 | 5 |
| 13 | 29 | 1 | 378 | 276 | 276 | 253 | 91 | 91 | 171 | 1,537 |
| 14 | 30 | 15 | 10 | 3 | 21 | 21 | 6 | 6 | 28 | 110 |
| 16 | 32 | 55 | 190 | 0 | 36 | 325 | 325 | 0 | 231 | 1,162 |
| 17 | 33 | 0 | 55 | 66 | 66 | 66 | 78 | 0 | 55 | 386 |
| 18 | 34 | 0 | 0 | 0 | 0 | 0 | 0 | 0 | 1 | 1 |
| 24 | 35 | 21 | 36 | 15 | 15 | 15 | 15 | 3 | 10 | 130 |
| 26 | 36 | 0 | 15 | 78 | 66 | 45 | 36 | 45 | 78 | 363 |
| 27 | 37 | 1 | 15 | 1 | 6 | 1 | 15 | 6 | 28 | 73 |
| Sum | 15 | 559 | 3,479 | 2,314 | 5,337 | 4,870 | 2,660 | 316 | 5,831 | 25,366 |

**Supplemental_Table_S9.** QTLR high LD blocks found in the eight lines, ordered by location of the first marker in the block. Chr - chromosome; Q - QTLR serial number (Supplemental_Table_S1); B - serial number of the high LD block within line (Supplemental_Table_S11); Start/End - location of the first and last SNP in the block; Length - size of the block (bp); Distance - base pairs between the first bp of the block and last bp of the block started before it on the same chromosome: coloured, negative distance indicates overlap of blocks (the block starts before the end of the previous block, ends after the start of the previous block); B OL - across lines number of blocks overlapping the current block; L OL - number of lines overlapping the current block.

| Line | C | Q | B | Start | End | Length | Distance | B OL | L OL |
|---|---|---|---|---|---|---|---|---|---|
| WPR1 | 1 | 3 | 1 | 52,276,776 | 52,277,046 | 271 | | 9 | 1 |
| WPR2 | 1 | 3 | 1 | 52,276,776 | 52,277,046 | 271 | -270 | 9 | 1 |
| WL2 | 1 | 3 | 1 | 52,276,776 | 52,637,288 | 360,513 | -270 | 20 | 1 |
| WPR1 | 1 | 3 | 2 | 52,276,779 | 52,276,917 | 139 | -360,509 | 8 | 1 |
| WPR2 | 1 | 3 | 2 | 52,276,779 | 52,276,917 | 139 | -138 | 8 | 1 |
| WL2 | 1 | 3 | 2 | 52,276,779 | 52,337,486 | 60,708 | -138 | 13 | 1 |
| WL1 | 1 | 3 | 1 | 52,276,779 | 52,408,209 | 131,431 | -60,707 | 15 | 1 |
| WL3 | 1 | 3 | 1 | 52,276,779 | 52,474,457 | 197,679 | -131,430 | 18 | 1 |
| RIR1 | 1 | 3 | 1 | 52,276,779 | 52,642,408 | 365,630 | -197,678 | 21 | 1 |
| WL2 | 1 | 3 | 3 | 52,276,940 | 52,642,408 | 365,469 | -365,468 | 19 | 1 |
| WL4 | 1 | 3 | 1 | 52,337,486 | 52,353,162 | 15,677 | -304,922 | 9 | 1 |
| WPR2 | 1 | 3 | 3 | 52,337,486 | 52,428,133 | 90,648 | -15,676 | 12 | 1 |
| WPR1 | 1 | 3 | 3 | 52,337,486 | 52,636,196 | 298,711 | -90,647 | 14 | 1 |
| WL3 | 1 | 3 | 2 | 52,337,486 | 52,642,408 | 304,923 | -298,710 | 17 | 1 |
| WL4 | 1 | 3 | 2 | 52,370,944 | 52,382,586 | 11,643 | -271,464 | 9 | 1 |
| WPR1 | 1 | 3 | 4 | 52,370,944 | 52,642,408 | 271,465 | -11,642 | 15 | 1 |
| WL4 | 1 | 3 | 3 | 52,428,133 | 52,474,457 | 46,325 | -214,275 | 10 | 1 |
| WL2 | 1 | 3 | 4 | 52,453,441 | 52,637,225 | 183,785 | -21,016 | 11 | 1 |
| WPR2 | 1 | 3 | 4 | 52,453,441 | 52,637,288 | 183,848 | -183,784 | 11 | 1 |
| WPR2 | 1 | 3 | 5 | 52,636,296 | 52,642,408 | 6,113 | -992 | 9 | 1 |
| WPR1 | 1 | 3 | 5 | 52,636,852 | 52,637,288 | 437 | -5,556 | 8 | 1 |
| WL4 | 1 | 3 | 4 | 52,641,813 | 52,642,408 | 596 | 4,525 | 5 | |
| WPR1 | 1 | 4 | 6 | 72,307,283 | 72,307,630 | 348 | 19,664,875 | 4 | 1 |
| WPR2 | 1 | 4 | 6 | 72,307,283 | 72,307,630 | 348 | -347 | 4 | 1 |
| RIR1 | 1 | 4 | 2 | 72,307,283 | 72,307,630 | 348 | -347 | 4 | 1 |
| WL3 | 1 | 4 | 3 | 72,307,291 | 72,307,616 | 326 | -339 | 4 | 1 |

| Line | C | Q | B | Start | End | Length | Distance | B OL | L OL |
|---|---|---|---|---|---|---|---|---|---|
| WL4 | 1 | 4 | 5 | 72,307,291 | 72,307,616 | 326 | -325 | 4 | 1 |
| WL3 | 1 | 5 | 4 | 77,440,144 | 77,864,774 | 424,631 | 5,132,528 | 13 | 1 |
| WL4 | 1 | 5 | 6 | 77,440,144 | 77,864,774 | 424,631 | -424,630 | 13 | 1 |
| WL2 | 1 | 5 | 5 | 77,440,155 | 77,440,159 | 5 | -424,619 | 3 | 1 |
| RIR1 | 1 | 5 | 3 | 77,440,155 | 78,054,406 | 614,252 | -4 | 20 | 1 |
| WPR1 | 1 | 5 | 7 | 77,442,145 | 77,444,931 | 2,787 | -612,261 | 5 | 1 |
| WL1 | 1 | 5 | 2 | 77,442,831 | 77,567,384 | 124,554 | -2,100 | 5 | 1 |
| WL2 | 1 | 5 | 6 | 77,442,831 | 77,567,384 | 124,554 | -124,553 | 5 | 1 |
| WPR1 | 1 | 5 | 8 | 77,862,924 | 77,863,325 | 402 | 295,540 | 7 | 1 |
| WL2 | 1 | 5 | 7 | 77,862,924 | 77,864,774 | 1,851 | -401 | 9 | 1 |
| RIR1 | 1 | 5 | 4 | 77,862,924 | 78,055,130 | 192,207 | -1,850 | 16 | 1 |
| WL1 | 1 | 5 | 3 | 77,863,325 | 77,864,774 | 1,450 | -191,805 | 9 | 1 |
| WPR2 | 1 | 5 | 7 | 77,863,325 | 78,051,237 | 187,913 | -1,449 | 16 | 1 |
| WPR2 | 1 | 5 | 8 | 77,863,508 | 78,051,418 | 187,911 | -187,729 | 15 | 1 |
| WPR1 | 1 | 5 | 9 | 77,864,774 | 78,051,418 | 186,645 | -186,644 | 15 | 1 |
| WL2 | 1 | 5 | 8 | 78,045,732 | 78,048,417 | 2,686 | -5,686 | 11 | 1 |
| WPR1 | 1 | 5 | 10 | 78,045,732 | 78,053,783 | 8,052 | -2,685 | 11 | 1 |
| WPR2 | 1 | 5 | 9 | 78,045,732 | 78,053,783 | 8,052 | -8,051 | 11 | 1 |
| RIR1 | 1 | 5 | 5 | 78,045,732 | 78,053,783 | 8,052 | -8,051 | 11 | 1 |
| WL3 | 1 | 5 | 5 | 78,045,732 | 78,054,406 | 8,675 | -8,051 | 11 | 1 |
| WL4 | 1 | 5 | 7 | 78,045,732 | 78,054,406 | 8,675 | -8,674 | 11 | 1 |
| WPR2 | 1 | 5 | 10 | 78,048,417 | 78,048,418 | 2 | -5,989 | 11 | 1 |
| WL2 | 1 | 7 | 9 | 104,379,743 | 104,392,252 | 12,510 | 26,331,325 | 9 | 1 |
| WL3 | 1 | 7 | 6 | 104,379,743 | 104,392,252 | 12,510 | -12,509 | 9 | 1 |
| WL4 | 1 | 7 | 8 | 104,379,743 | 104,392,252 | 12,510 | -12,509 | 9 | 1 |
| WPR1 | 1 | 7 | 11 | 104,379,743 | 104,418,467 | 38,725 | -12,509 | 9 | 1 |
| WPR2 | 1 | 7 | 11 | 104,379,743 | 104,418,467 | 38,725 | -38,724 | 9 | 1 |
| WL3 | 1 | 7 | 7 | 104,383,567 | 104,385,136 | 1,570 | -34,900 | 9 | 1 |
| WL1 | 1 | 7 | 4 | 104,383,567 | 104,387,123 | 3,557 | -1,569 | 9 | 1 |
| WPR2 | 1 | 7 | 12 | 104,383,567 | 104,387,123 | 3,557 | -3,556 | 9 | 1 |
| WL4 | 1 | 7 | 9 | 104,383,654 | 104,385,136 | 1,483 | -3,469 | 9 | 1 |
| RIR1 | 1 | 7 | 6 | 104,383,654 | 104,387,123 | 3,470 | -1,482 | 9 | 1 |
| WL2 | 1 | 7 | 10 | 106,098,899 | 106,098,942 | 44 | 1,711,776 | 2 | 1 |
| WL3 | 1 | 7 | 8 | 106,098,899 | 106,098,942 | 44 | -43 | 2 | 1 |
| WL4 | 1 | 7 | 10 | 106,098,899 | 106,098,942 | 44 | -43 | 2 | 1 |
| RIR1 | 1 | 9 | 7 | 173,657,608 | 173,682,298 | 24,691 | 67,558,666 | 14 | 1 |
| WPR2 | 1 | 9 | 13 | 173,657,608 | 173,691,228 | 33,621 | -24,690 | 16 | 1 |
| RIR1 | 1 | 9 | 8 | 173,658,086 | 173,664,538 | 6,453 | -33,142 | 10 | 1 |
| WPR1 | 1 | 9 | 12 | 173,658,086 | 173,686,475 | 28,390 | -6,452 | 14 | 1 |
| WL1 | 1 | 9 | 5 | 173,658,646 | 173,662,588 | 3,943 | -27,829 | 10 | 1 |
| WL2 | 1 | 9 | 11 | 173,658,646 | 173,662,588 | 3,943 | -3,942 | 10 | 1 |
| WL3 | 1 | 9 | 9 | 173,658,646 | 173,662,588 | 3,943 | -3,942 | 10 | 1 |
| WL5 | 1 | 9 | 1 | 173,658,646 | 173,662,588 | 3,943 | -3,942 | 10 | 1 |
| RIR1 | 1 | 9 | 9 | 173,658,646 | 173,670,185 | 11,540 | -3,942 | 12 | 1 |

| Line | C | Q | B | Start | End | Length | Distance | B OL | L OL |
|---|---|---|---|---|---|---|---|---|---|
| WL4 | 1 | 9 | 11 | 173,660,604 | 173,662,588 | 1,985 | -9,581 | 10 | 1 |
| RIR1 | 1 | 9 | 10 | 173,660,604 | 173,691,545 | 30,942 | -1,984 | 16 | 1 |
| RIR1 | 1 | 9 | 11 | 173,667,742 | 173,691,228 | 23,487 | -23,803 | 10 | 1 |
| WL4 | 1 | 9 | 12 | 173,670,185 | 173,681,300 | 11,116 | -21,043 | 8 | 1 |
| RIR1 | 1 | 9 | 12 | 173,681,300 | 173,684,220 | 2,921 | 0 | 7 | 1 |
| WL2 | 1 | 9 | 12 | 173,681,300 | 173,691,545 | 10,246 | -2,920 | 9 | 1 |
| WL1 | 1 | 9 | 6 | 173,691,228 | 173,691,545 | 318 | -317 | 5 | 1 |
| WL4 | 1 | 9 | 13 | 173,691,228 | 177,386,285 | 3,695,058 | -317 | 18 | 1 |
| WPR2 | 1 | 10 | 14 | 177,365,684 | 177,365,770 | 87 | -20,601 | 6 | 1 |
| WL4 | 1 | 10 | 14 | 177,365,684 | 177,365,770 | 87 | -86 | 6 | 1 |
| RIR1 | 1 | 1 | 13 | 177,365,684 | 177,366,081 | 398 | -86 | 8 | 1 |
| WL2 | 1 | 10 | 13 | 177,365,684 | 177,369,603 | 3,920 | -397 | 9 | 1 |
| WL3 | 1 | 10 | 10 | 177,365,684 | 177,369,603 | 3,920 | -3,919 | 9 | 1 |
| WL5 | 1 | 10 | 2 | 177,365,684 | 177,369,603 | 3,920 | -3,919 | 9 | 1 |
| WPR1 | 1 | 10 | 13 | 177,366,081 | 177,387,725 | 21,645 | -3,522 | 11 | 1 |
| WPR2 | 1 | 10 | 15 | 177,366,081 | 177,387,725 | 21,645 | -21,644 | 11 | 1 |
| RIR1 | 1 | 1 | 14 | 177,369,603 | 177,389,132 | 19,530 | -18,122 | 10 | 1 |
| WL2 | 1 | 10 | 14 | 177,385,869 | 177,386,295 | 427 | -3,263 | 7 | 1 |
| RIR1 | 1 | 1 | 15 | 177,385,869 | 177,386,295 | 427 | -426 | 7 | 1 |
| WL3 | 1 | 10 | 11 | 177,385,869 | 177,388,614 | 2,746 | -426 | 7 | 1 |
| WL4 | 1 | 10 | 15 | 177,385,869 | 177,388,614 | 2,746 | -2,745 | 7 | 1 |
| WPR1 | 1 | 11 | 14 | 196,386,357 | 196,389,592 | 3,236 | 18,997,743 | 8 | 1 |
| WL4 | 1 | 11 | 16 | 196,386,494 | 196,386,900 | 407 | -3,098 | 8 | 1 |
| WL1 | 1 | 11 | 7 | 196,386,494 | 196,390,599 | 4,106 | -406 | 8 | 1 |
| WL2 | 1 | 11 | 15 | 196,386,494 | 196,390,599 | 4,106 | -4,105 | 8 | 1 |
| WL5 | 1 | 11 | 3 | 196,386,494 | 196,395,140 | 8,647 | -4,105 | 8 | 1 |
| RIR1 | 1 | 11 | 16 | 196,386,900 | 196,389,592 | 2,693 | -8,240 | 8 | 1 |
| WL3 | 1 | 11 | 12 | 196,386,900 | 196,389,598 | 2,699 | -2,692 | 8 | 1 |
| WPR2 | 1 | 11 | 16 | 196,386,900 | 196,389,598 | 2,699 | -2,698 | 8 | 1 |
| WPR1 | 1 | 11 | 15 | 196,386,900 | 196,395,140 | 8,241 | -2,698 | 8 | 1 |
| RIR1 | 2 | 12 | 1 | 633,591 | 633,592 | 2 |  | 0 |  |
| WPR1 | 2 | 13 | 1 | 45,900,042 | 45,921,445 | 21,404 | 45,266,450 | 13 | 1 |
| WPR2 | 2 | 13 | 1 | 45,900,042 | 45,921,445 | 21,404 | -21,403 | 13 | 1 |
| RIR1 | 2 | 13 | 2 | 45,900,042 | 45,932,906 | 32,865 | -21,403 | 13 | 1 |
| WL5 | 2 | 13 | 1 | 45,900,106 | 45,902,476 | 2,371 | -32,800 | 8 | 1 |
| WL3 | 2 | 13 | 1 | 45,900,106 | 45,918,845 | 18,740 | -2,370 | 13 | 1 |
| WL1 | 2 | 12 | 1 | 45,900,106 | 45,935,931 | 35,826 | -18,739 | 13 | 1 |
| WL1 | 2 | 13 | 2 | 45,901,252 | 45,902,086 | 835 | -34,679 | 8 | 1 |
| WL2 | 2 | 12 | 1 | 45,901,252 | 45,918,845 | 17,594 | -834 | 13 | 1 |
| RIR1 | 2 | 13 | 3 | 45,902,086 | 45,922,309 | 20,224 | -16,759 | 13 | 1 |
| WL2 | 2 | 13 | 2 | 45,911,199 | 45,935,931 | 24,733 | -11,110 | 11 | 1 |
| WL3 | 2 | 13 | 2 | 45,911,199 | 45,935,931 | 24,733 | -24,732 | 11 | 1 |
| WPR1 | 2 | 13 | 2 | 45,917,333 | 45,932,906 | 15,574 | -18,598 | 11 | 1 |
| WPR2 | 2 | 13 | 2 | 45,917,333 | 45,932,906 | 15,574 | -15,573 | 11 | 1 |

| Line | C | Q | B | Start | End | Length | Distance | B OL | L OL |
|---|---|---|---|---|---|---|---|---|---|
| RIR1 | 2 | 13 | 4 | 45,917,978 | 45,927,742 | 9,765 | -14,928 | 11 | 1 |
| WL2 | 4 | 19 | 1 | 87,451,251 | 87,483,115 | 31,865 | | 9 | 1 |
| WPR2 | 4 | 19 | 1 | 87,451,251 | 87,535,920 | 84,670 | -31,864 | 9 | 1 |
| WL2 | 4 | 19 | 2 | 87,455,417 | 87,461,520 | 6,104 | -80,503 | 5 | 1 |
| WL1 | 4 | 19 | 1 | 87,455,417 | 87,469,358 | 13,942 | -6,103 | 5 | 1 |
| WL3 | 4 | 19 | 1 | 87,455,417 | 87,469,358 | 13,942 | -13,941 | 5 | 1 |
| WL4 | 4 | 19 | 1 | 87,455,417 | 87,469,358 | 13,942 | -13,941 | 5 | 1 |
| WL2 | 4 | 19 | 3 | 87,479,459 | 87,482,774 | 3,316 | 10,101 | 5 | 1 |
| WL3 | 4 | 19 | 2 | 87,479,459 | 87,482,774 | 3,316 | -3,315 | 5 | 1 |
| WPR2 | 4 | 19 | 2 | 87,479,459 | 87,482,774 | 3,316 | -3,315 | 5 | 1 |
| WPR1 | 4 | 19 | 1 | 87,479,459 | 87,535,920 | 56,462 | -3,315 | 5 | 1 |
| WL2 | 5 | 20 | 1 | 8,421,567 | 8,422,079 | 513 | | 2 | 1 |
| WL1 | 5 | 20 | 1 | 8,422,079 | 8,777,825 | 355,747 | 0 | 7 | 1 |
| RIR1 | 5 | 20 | 1 | 8,422,079 | 8,778,394 | 356,316 | -355,746 | 7 | 1 |
| WL5 | 5 | 20 | 1 | 8,544,549 | 8,777,825 | 233,277 | -233,845 | 6 | 1 |
| WPR1 | 5 | 20 | 1 | 8,544,933 | 8,545,143 | 211 | -232,892 | 5 | 1 |
| WPR2 | 5 | 20 | 1 | 8,544,933 | 8,545,143 | 211 | -210 | 5 | 1 |
| WL2 | 5 | 20 | 2 | 8,545,143 | 8,777,825 | 232,683 | 0 | 6 | 1 |
| WPR2 | 5 | 20 | 2 | 8,777,825 | 8,778,679 | 855 | 0 | 4 | 1 |
| WL4 | 5 | 21 | 1 | 19,756,582 | 19,803,706 | 47,125 | 10,977,903 | 7 | 1 |
| WL5 | 5 | 21 | 2 | 19,756,582 | 20,707,750 | 951,169 | -47,124 | 21 | 1 |
| WL4 | 5 | 21 | 2 | 19,756,660 | 19,757,145 | 486 | -951,090 | 6 | 1 |
| WL1 | 5 | 21 | 2 | 19,756,660 | 19,803,706 | 47,047 | -485 | 7 | 1 |
| WPR2 | 5 | 21 | 3 | 19,756,660 | 19,803,706 | 47,047 | -47,046 | 7 | 1 |
| RIR1 | 5 | 21 | 2 | 19,756,660 | 20,723,673 | 967,014 | -47,046 | 23 | 1 |
| WL3 | 5 | 21 | 1 | 19,756,660 | 20,733,573 | 976,914 | -967,013 | 23 | 1 |
| WL2 | 5 | 21 | 3 | 19,803,706 | 19,804,039 | 334 | -929,867 | 7 | 1 |
| WL4 | 5 | 21 | 3 | 19,804,039 | 20,723,673 | 919,635 | 0 | 19 | 1 |
| WPR1 | 5 | 21 | 2 | 20,580,772 | 20,589,955 | 9,184 | -142,901 | 9 | 1 |
| WPR2 | 5 | 21 | 4 | 20,580,772 | 20,589,955 | 9,184 | -9,183 | 9 | 1 |
| WL4 | 5 | 21 | 4 | 20,580,772 | 20,589,955 | 9,184 | -9,183 | 9 | 1 |
| WL5 | 5 | 21 | 3 | 20,580,772 | 20,596,531 | 15,760 | -9,183 | 12 | 1 |
| WL2 | 5 | 21 | 4 | 20,580,772 | 20,628,840 | 48,069 | -15,759 | 14 | 1 |
| RIR1 | 5 | 21 | 3 | 20,589,955 | 20,733,573 | 143,619 | -38,885 | 18 | 1 |
| WL4 | 5 | 21 | 5 | 20,596,531 | 20,628,840 | 32,310 | -137,042 | 11 | 1 |
| WPR1 | 5 | 21 | 3 | 20,596,531 | 20,723,673 | 127,143 | -32,309 | 15 | 1 |
| WPR2 | 5 | 21 | 5 | 20,596,531 | 20,723,673 | 127,143 | -127,142 | 15 | 1 |
| WPR1 | 5 | 21 | 4 | 20,612,350 | 20,733,573 | 121,224 | -111,323 | 14 | 1 |
| WPR2 | 5 | 21 | 6 | 20,612,350 | 20,733,573 | 121,224 | -121,223 | 14 | 1 |
| WL2 | 5 | 21 | 5 | 20,665,999 | 20,723,673 | 57,675 | -67,574 | 12 | 1 |
| WL5 | 5 | 21 | 4 | 20,672,487 | 20,723,673 | 51,187 | -51,186 | 12 | 1 |
| WL4 | 5 | 21 | 6 | 20,716,370 | 20,733,573 | 17,204 | -7,303 | 11 | 1 |
| WL5 | 5 | 21 | 5 | 20,716,370 | 20,733,573 | 17,204 | -17,203 | 11 | 1 |
| WL4 | 6 | 24 | 1 | 32,573,870 | 32,590,409 | 16,540 | | 8 | 1 |

| Line | C | Q | B | Start | End | Length | Distance | B OL | L OL |
|---|---|---|---|---|---|---|---|---|---|
| WPR1 | 6 | 24 | 1 | 32,573,870 | 32,713,410 | 139,541 | -16,539 | 19 | 1 |
| WPR2 | 6 | 24 | 1 | 32,573,870 | 32,713,410 | 139,541 | -139,540 | 19 | 1 |
| WL1 | 6 | 24 | 1 | 32,573,870 | 32,803,638 | 229,769 | -139,540 | 19 | 1 |
| WL3 | 6 | 24 | 1 | 32,573,870 | 32,803,638 | 229,769 | -229,768 | 19 | 1 |
| WL2 | 6 | 24 | 1 | 32,579,493 | 32,588,194 | 8,702 | -224,145 | 8 | 1 |
| WL4 | 6 | 24 | 2 | 32,579,493 | 32,803,462 | 223,970 | -8,701 | 19 | 1 |
| WL4 | 6 | 24 | 2 | 32,583,416 | 32,635,144 | 51,729 | -220,046 | 13 | 1 |
| WL5 | 6 | 24 | 1 | 32,583,416 | 32,635,255 | 51,840 | -51,728 | 15 | 1 |
| WPR2 | 6 | 24 | 2 | 32,634,574 | 32,635,144 | 571 | -681 | 11 | 1 |
| WPR1 | 6 | 24 | 2 | 32,634,574 | 32,644,318 | 9,745 | -570 | 14 | 1 |
| RIR1 | 6 | 24 | 1 | 32,634,574 | 32,644,318 | 9,745 | -9,744 | 14 | 1 |
| WL2 | 6 | 24 | 2 | 32,634,574 | 32,713,410 | 78,837 | -9,744 | 17 | 1 |
| WL2 | 6 | 24 | 3 | 32,635,144 | 32,642,143 | 7,000 | -78,266 | 14 | 1 |
| WL5 | 6 | 24 | 3 | 32,635,255 | 32,642,143 | 6,889 | -6,888 | 12 | 1 |
| WL4 | 6 | 24 | 3 | 32,635,255 | 32,803,638 | 168,384 | -6,888 | 15 | 1 |
| RIR1 | 6 | 24 | 2 | 32,642,143 | 32,698,617 | 56,475 | -161,495 | 12 | 1 |
| WL2 | 6 | 24 | 4 | 32,698,298 | 32,712,118 | 13,821 | -319 | 9 | 1 |
| WL4 | 6 | 24 | 4 | 32,712,118 | 32,713,410 | 1,293 | 0 | 9 | 1 |
| RIR1 | 6 | 24 | 3 | 32,713,410 | 32,803,638 | 90,229 | 0 | 8 | 1 |
| WL5 | 7 | 25 | 1 | 13,676,441 | 16,772,561 | 3,096,121 |  | 0 |  |
| WL3 | 13 | 29 | 1 | 12,164,061 | 12,170,453 | 6,393 |  | 17 | 1 |
| WPR2 | 13 | 29 | 1 | 12,164,061 | 12,171,514 | 7,454 | -6,392 | 19 | 1 |
| WPR1 | 13 | 29 | 1 | 12,164,061 | 12,188,510 | 24,450 | -7,453 | 21 | 1 |
| RIR1 | 13 | 29 | 1 | 12,164,061 | 12,188,510 | 24,450 | -24,449 | 21 | 1 |
| WL2 | 13 | 29 | 1 | 12,164,859 | 12,171,514 | 6,656 | -23,651 | 19 | 1 |
| WPR1 | 13 | 29 | 2 | 12,164,859 | 12,182,209 | 17,351 | -6,655 | 21 | 1 |
| WPR2 | 13 | 29 | 2 | 12,164,859 | 12,182,209 | 17,351 | -17,350 | 21 | 1 |
| WL3 | 13 | 29 | 2 | 12,164,859 | 12,187,242 | 22,384 | -17,350 | 21 | 1 |
| WL4 | 13 | 29 | 1 | 12,164,859 | 12,188,510 | 23,652 | -22,383 | 21 | 1 |
| WL3 | 13 | 29 | 3 | 12,165,490 | 12,188,510 | 23,021 | -23,020 | 21 | 1 |
| WL2 | 13 | 29 | 2 | 12,166,511 | 12,173,136 | 6,626 | -21,999 | 20 | 1 |
| WPR1 | 13 | 29 | 3 | 12,166,511 | 12,187,242 | 20,732 | -6,625 | 21 | 1 |
| WPR2 | 13 | 29 | 3 | 12,166,511 | 12,187,242 | 20,732 | -20,731 | 21 | 1 |
| WL5 | 13 | 29 | 1 | 12,166,511 | 12,188,510 | 22,000 | -20,731 | 21 | 1 |
| WL2 | 13 | 29 | 3 | 12,167,802 | 12,168,514 | 713 | -20,708 | 16 | 1 |
| WPR1 | 13 | 29 | 4 | 12,167,802 | 12,170,453 | 2,652 | -712 | 17 | 1 |
| RIR1 | 13 | 29 | 2 | 12,167,802 | 12,182,246 | 14,445 | -2,651 | 21 | 1 |
| RIR1 | 13 | 29 | 3 | 12,170,453 | 12,186,115 | 15,663 | -11,793 | 20 | 1 |
| WL2 | 13 | 29 | 4 | 12,170,561 | 12,174,479 | 3,919 | -15,554 | 17 | 1 |
| WL2 | 13 | 29 | 5 | 12,171,402 | 12,188,510 | 17,109 | -3,077 | 18 |  |
| WL2 | 13 | 29 | 6 | 12,172,027 | 12,187,242 | 15,216 | -16,483 | 16 |  |
| WL2 | 13 | 29 | 7 | 12,182,052 | 12,182,209 | 158 | -5,190 | 14 |  |
| WL5 | 14 | 30 | 1 | 9,343,521 | 9,344,120 | 600 |  | 3 | 1 |
| WPR1 | 14 | 30 | 1 | 9,343,654 | 9,344,865 | 1,212 | -466 | 3 | 1 |

| Line | C | Q | B | Start | End | Length | Distance | B OL | L OL |
|---|---|---|---|---|---|---|---|---|---|
| WPR2 | 14 | 30 | 1 | 9,343,654 | 9,344,865 | 1,212 | -1,211 | 3 | 1 |
| RIR1 | 14 | 30 | 1 | 9,343,654 | 9,344,865 | 1,212 | -1,211 | 3 | 1 |
| WL4 | 16 | 32 | 1 | 2,357,192 | 2,473,550 | 116,359 | | 3 | 1 |
| RIR1 | 16 | 32 | 1 | 2,357,192 | 2,551,587 | 194,396 | -116,358 | 15 | 1 |
| WL2 | 16 | 32 | 1 | 2,357,192 | 2,598,372 | 241,181 | -194,395 | 16 | 1 |
| WPR2 | 16 | 32 | 1 | 2,473,550 | 2,597,942 | 124,393 | -124,822 | 15 | 1 |
| WPR2 | 16 | 32 | 2 | 2,475,052 | 2,476,811 | 1,760 | -122,890 | 8 | 1 |
| RIR1 | 16 | 32 | 2 | 2,475,052 | 2,600,004 | 124,953 | -1,759 | 15 | 1 |
| WL2 | 16 | 32 | 2 | 2,475,052 | 2,600,170 | 125,119 | -124,952 | 16 | 1 |
| WL2 | 16 | 32 | 3 | 2,476,123 | 2,599,123 | 123,001 | -124,047 | 15 | 1 |
| WL1 | 16 | 32 | 1 | 2,476,467 | 2,600,920 | 124,454 | -122,656 | 16 | 1 |
| WL4 | 16 | 32 | 2 | 2,476,811 | 2,476,874 | 64 | -124,109 | 9 | 1 |
| RIR1 | 16 | 32 | 3 | 2,476,874 | 2,600,920 | 124,047 | 0 | 15 | 1 |
| WPR2 | 16 | 32 | 3 | 2,477,464 | 2,477,515 | 52 | -123,456 | 9 | 1 |
| WL4 | 16 | 32 | 3 | 2,477,464 | 2,477,515 | 52 | -51 | 9 | 1 |
| WL4 | 16 | 32 | 4 | 2,550,271 | 2,597,967 | 47,697 | 72,756 | 11 | 1 |
| WPR1 | 16 | 32 | 1 | 2,550,271 | 2,600,004 | 49,734 | -47,696 | 11 | 1 |
| WPR2 | 16 | 32 | 4 | 2,551,587 | 2,600,004 | 48,418 | -48,417 | 11 | 1 |
| WPR2 | 16 | 32 | 5 | 2,597,967 | 2,599,123 | 1,157 | -2,037 | 9 | |
| WPR2 | 16 | 32 | 6 | 2,600,170 | 2,600,920 | 751 | 1,047 | 3 | |
| WL3 | 17 | 33 | 1 | 3,938,971 | 3,940,192 | 1,222 | | 6 | 1 |
| WL4 | 17 | 33 | 1 | 3,938,971 | 3,940,192 | 1,222 | -1,221 | 6 | 1 |
| WPR2 | 17 | 33 | 1 | 3,938,971 | 3,941,957 | 2,987 | -1,221 | 10 | 1 |
| WPR1 | 17 | 33 | 1 | 3,938,971 | 3,944,711 | 5,741 | -2,986 | 14 | 1 |
| WL1 | 17 | 33 | 1 | 3,938,971 | 4,364,160 | 425,190 | -5,740 | 17 | 1 |
| WL2 | 17 | 33 | 1 | 3,938,971 | 4,364,160 | 425,190 | -425,189 | 17 | 1 |
| RIR1 | 17 | 33 | 1 | 3,940,192 | 3,941,957 | 1,766 | -423,968 | 10 | 1 |
| WL1 | 17 | 33 | 2 | 3,941,832 | 3,944,711 | 2,880 | -125 | 12 | 1 |
| WL2 | 17 | 33 | 2 | 3,941,832 | 3,944,711 | 2,880 | -2,879 | 12 | 1 |
| WL3 | 17 | 33 | 2 | 3,941,832 | 3,944,711 | 2,880 | -2,879 | 12 | 1 |
| WL4 | 17 | 33 | 2 | 3,941,832 | 3,944,711 | 2,880 | -2,879 | 12 | 1 |
| RIR1 | 17 | 33 | 2 | 3,942,076 | 3,943,548 | 1,473 | -2,635 | 10 | 1 |
| WL3 | 17 | 33 | 3 | 3,942,076 | 3,944,254 | 2,179 | -1,472 | 10 | 1 |
| WPR1 | 17 | 33 | 2 | 3,943,548 | 3,944,254 | 707 | -706 | 10 | 1 |
| WPR2 | 17 | 33 | 2 | 3,943,548 | 3,944,711 | 1,164 | -706 | 10 | 1 |
| WL4 | 17 | 33 | 3 | 4,358,764 | 4,364,103 | 5,340 | 414,053 | 3 | 1 |
| WL3 | 17 | 33 | 4 | 4,358,764 | 4,368,888 | 10,125 | -5,339 | 4 | 1 |
| WL4 | 17 | 33 | 4 | 4,364,160 | 4,368,888 | 4,729 | -4,728 | 3 | 1 |
| WL4 | 24 | 35 | 1 | 4,813,725 | 4,813,744 | 20 | | 2 | 1 |
| WL1 | 24 | 35 | 1 | 4,813,725 | 4,813,815 | 91 | -19 | 2 | 1 |
| WL2 | 24 | 35 | 1 | 4,813,725 | 4,813,815 | 91 | -90 | 2 | 1 |
| WPR2 | 24 | 35 | 1 | 5,170,427 | 5,238,859 | 68,433 | 356,612 | 7 | 1 |
| WL2 | 24 | 35 | 2 | 5,237,664 | 5,238,307 | 644 | -1,195 | 7 | 1 |
| WL5 | 24 | 35 | 1 | 5,237,664 | 5,238,724 | 1,061 | -643 | 7 | 1 |

| Line | C | Q | B | Start | End | Length | Distance | B OL | L OL |
|---|---|---|---|---|---|---|---|---|---|
| WL1 | 24 | 35 | 2 | 5,237,664 | 5,238,833 | 1,170 | -1,060 | 7 | 1 |
| WL3 | 24 | 35 | 1 | 5,237,664 | 5,238,833 | 1,170 | -1,169 | 7 | 1 |
| WL4 | 24 | 35 | 2 | 5,237,664 | 5,238,833 | 1,170 | -1,169 | 7 | 1 |
| WPR1 | 24 | 35 | 1 | 5,238,307 | 5,238,859 | 553 | -526 | 7 | 1 |
| RIR1 | 24 | 35 | 1 | 5,238,307 | 5,238,859 | 553 | -552 | 7 | 1 |
| WL2 | 26 | 36 | 1 | 4,691,657 | 4,759,369 | 67,713 |  | 11 | 1 |
| RIR1 | 26 | 36 | 1 | 4,691,657 | 4,759,889 | 68,233 | -67,712 | 12 | 1 |
| WL3 | 26 | 36 | 1 | 4,691,657 | 4,759,960 | 68,304 | -68,232 | 12 | 1 |
| WPR1 | 26 | 36 | 1 | 4,691,657 | 4,759,960 | 68,304 | -68,303 | 12 | 1 |
| WPR2 | 26 | 36 | 1 | 4,691,657 | 4,759,960 | 68,304 | -68,303 | 12 | 1 |
| WL5 | 26 | 36 | 1 | 4,691,657 | 4,759,960 | 68,304 | -68,303 | 12 | 1 |
| WL4 | 26 | 36 | 1 | 4,691,657 | 4,761,708 | 70,052 | -68,303 | 14 | 1 |
| WL3 | 26 | 36 | 2 | 4,755,292 | 4,755,531 | 240 | -6,416 | 9 | 1 |
| RIR1 | 26 | 36 | 2 | 4,755,292 | 4,761,708 | 6,417 | -239 | 14 | 1 |
| RIR1 | 26 | 36 | 3 | 4,755,531 | 4,759,960 | 4,430 | -6,177 | 12 | 1 |
| WL3 | 26 | 36 | 3 | 4,756,562 | 4,759,369 | 2,808 | -3,398 | 10 | 1 |
| RIR1 | 26 | 36 | 4 | 4,756,606 | 4,756,618 | 13 | -2,763 | 10 | 1 |
| WL5 | 26 | 36 | 2 | 4,759,648 | 4,761,708 | 2,061 | 3,030 | 10 | 1 |
| WL3 | 26 | 36 | 4 | 4,761,468 | 4,761,708 | 241 | -240 | 4 | 1 |
| WPR1 | 26 | 36 | 2 | 4,761,468 | 4,761,708 | 241 | -240 | 4 | 1 |
| WL2 | 27 | 37 | 1 | 3,923,054 | 3,923,089 | 36 |  | 4 | 1 |
| WPR1 | 27 | 37 | 1 | 3,923,054 | 3,923,089 | 36 | -35 | 4 | 1 |
| WL4 | 27 | 37 | 1 | 3,923,054 | 3,923,089 | 36 | -35 | 4 | 1 |
| WL5 | 27 | 37 | 1 | 3,923,054 | 3,923,089 | 36 | -35 | 4 | 1 |
| RIR1 | 27 | 37 | 1 | 3,923,054 | 3,923,089 | 36 | -35 | 4 | 1 |
| RIR1 | 27 | 37 | 2 | 4,040,797 | 4,043,381 | 2,585 | 117,708 | 3 | 1 |
| WL1 | 27 | 37 | 1 | 4,040,797 | 4,064,529 | 23,733 | -2,584 | 3 | 1 |
| WL4 | 27 | 37 | 2 | 4,040,797 | 4,064,529 | 23,733 | -23,732 | 3 | 1 |
| WL2 | 27 | 37 | 2 | 4,043,381 | 4,045,076 | 1,696 | -21,148 | 3 | 1 |

74

**Supplemental_Table_S10.** Distribution of the lengths of high LD blocks in the $F_6$ QTLRs tested in the eight pure lines, pooled together. Length - size of an LD block in bp (note the different scales of the three parts of the table, separated by thick lines); Blocks - number of blocks; % - proportion of the LD blocks among all blocks.

| Length | Blocks | % |
|---:|---:|---:|
| ≤10,000 | 141 | 0.513 |
| >10,000 - 20,000 | 28 | 0.102 |
| >20,000 - 30,000 | 20 | 0.073 |
| >30,000 - 40,000 | 8 | 0.029 |
| >40,000 - 50,000 | 8 | 0.029 |
| >50,000 - 100,000 | 19 | 0.069 |
| >100,000 - 150,000 | 17 | 0.062 |
| >150,000 - 200,000 | 9 | 0.033 |
| >200,000 - 250,000 | 6 | 0.022 |
| >250,000 - 300,000 | 2 | 0.007 |
| >300,000 - 350,000 | 1 | 0.004 |
| >350,000 - 400,000 | 5 | 0.018 |
| >400,000 - 450,000 | 4 | 0.015 |
| >450,000 - 500,000 | 0 | 0.000 |
| >500,000 - 1,000,000 | 5 | 0.018 |
| >1,000,000 - 1,500,000 | 0 | 0.000 |
| >1,500,000 - 2,000,000 | 0 | 0.000 |
| >2,000,000 - 2,500,000 | 0 | 0.000 |
| >2,500,000 - 3,000,000 | 0 | 0.000 |
| >3,000,000 - 3,500,000 | 1 | 0.004 |
| >3,500,000 - 4,000,000 | 1 | 0.004 |
| Sum | 275 | 1.000 |

**Supplemental_Table_S11.** QTLR high LD blocks found in the eight lines, aligned between lines, by location. C - chromosome, separated by thick lines; Q - QTLR serial number (Supplemental_Table_S1), colored by QTLR; bp - location of the marker in bp on the chicken GRCg6a reference; Distance - base pairs between the marker and the previous one; Marker - marker name, colored by the genomic element (some names of the markers include the genomic element name, e.g., lncRNA, long non-coding RNA; CSTA, the gene *CSTA*); Line blocks are numbered and colored independently within a line (that is, Block 1 in one line is not necessarily Block 1 in the other lines; same colour in different lines does not imply same block across lines); 0 - the marker had no high LD with any other marker; "-" - the marker was not informative in the line and thus was not tested in that line.

| C | Q | bp | Distance | Marker | Line blocks | | | | | | | |
|---|---|---|---|---|---|---|---|---|---|---|---|---|
| | | | | | WL1 | WL2 | WL3 | WPR1 | WPR2 | WL4 | WL5 | RIR1 |
| 1 | 3 | 52,276,776 | | lncRNA02.01 | - | 1 | - | 1 | 1 | - | - | - |
| 1 | 3 | 52,276,779 | 3 | lncRNA02.02 | 1 | 2 | 1 | 2 | 2 | 0 | - | 1 |
| 1 | 3 | 52,276,917 | 138 | lncRNA02.03 | 1 | 2 | 1 | 2 | 2 | 0 | - | 1 |
| 1 | 3 | 52,276,940 | 23 | lncRNA02.04 | 1 | 3 | 1 | 0 | 0 | 0 | - | 1 |
| 1 | 3 | 52,276,945 | 5 | lncRNA02.05 | - | 1 | - | 1 | 1 | - | - | - |
| 1 | 3 | 52,277,046 | 101 | lncRNA02.06 | - | 1 | - | 1 | 1 | - | - | - |
| 1 | 3 | 52,337,486 | 60,440 | lncRNA05.20 | 1 | 2 | 2 | 3 | 3 | 1 | - | 1 |
| 1 | 3 | 52,353,162 | 15,676 | lncRNA05.21 | 1 | 3 | 2 | - | - | 1 | - | 1 |
| 1 | 3 | 52,370,944 | 17,782 | lncRNA05.22 | 1 | 1 | - | 4 | 0 | 2 | - | 1 |
| 1 | 3 | 52,382,586 | 11,642 | lncRNA05.23 | - | - | 2 | 0 | 0 | 2 | - | 1 |
| 1 | 3 | 52,400,103 | 17,517 | lncRNA05.24 | - | 0 | 1 | - | - | - | - | - |
| 1 | 3 | 52,408,209 | 8,106 | lncRNA05.25 | 1 | 1 | 2 | 3 | 3 | - | - | - |
| 1 | 3 | 52,428,133 | 19,924 | lncRNA05.26 | - | 0 | 2 | 3 | 3 | 3 | - | 1 |
| 1 | 3 | 52,453,441 | 25,308 | lncRNA05.27 | - | 4 | - | 0 | 4 | - | - | 1 |
| 1 | 3 | 52,468,434 | 14,993 | lncRNA05.28 | - | 3 | 1 | 4 | 0 | 3 | - | - |
| 1 | 3 | 52,474,457 | 6,023 | lncRNA05.29 | - | 3 | 1 | - | - | 3 | - | - |
| 1 | 3 | 52,636,196 | 161,739 | lncRNA03.07 | - | - | - | 3 | 0 | - | - | - |
| 1 | 3 | 52,636,296 | 100 | lncRNA03.08 | - | - | - | 4 | 5 | - | - | - |
| 1 | 3 | 52,636,440 | 144 | lncRNA03.09 | - | - | - | 4 | 5 | - | - | - |
| 1 | 3 | 52,636,627 | 187 | lncRNA03.10 | - | - | - | - | - | - | - | 1 |
| 1 | 3 | 52,636,852 | 225 | lncRNA03.11 | - | 1 | - | 5 | 4 | - | - | 1 |
| 1 | 3 | 52,637,175 | 323 | lncRNA03.12 | - | 1 | - | 5 | 4 | - | - | 1 |
| 1 | 3 | 52,637,225 | 50 | lncRNA03.13 | - | 4 | - | 4 | 5 | - | - | - |
| 1 | 3 | 52,637,288 | 63 | lncRNA03.14 | - | 1 | - | 5 | 4 | - | - | 1 |
| 1 | 3 | 52,638,688 | 1,400 | lncRNA04.15 | - | - | - | 4 | 5 | - | - | - |
| 1 | 3 | 52,641,813 | 3,125 | lncRNA04.16 | - | 3 | 2 | 4 | 5 | 4 | - | 1 |
| 1 | 3 | 52,641,890 | 77 | lncRNA04.17 | - | 3 | 2 | 4 | 5 | 4 | - | 1 |
| 1 | 3 | 52,642,224 | 334 | lncRNA04.18 | - | 3 | 2 | - | - | 4 | - | 1 |

| C | Q | bp | Distance | Marker | Line blocks | | | | | | | |
|---|---|---|---|---|---|---|---|---|---|---|---|---|
| | | | | | WL1 | WL2 | WL3 | WPR1 | WPR2 | WL4 | WL5 | RIR1 |
| 1 | 3 | 52,642,408 | 184 | lncRNA04.19 | - | 3 | 2 | 4 | 5 | 4 | - | 1 |
| 1 | 4 | 72,307,283 | 19,664,875 | lncRNA01.01 | - | - | - | 6 | 6 | - | - | 2 |
| 1 | 4 | 72,307,291 | 8 | lncRNA01.02 | - | - | 3 | - | - | 5 | - | - |
| 1 | 4 | 72,307,369 | 78 | lncRNA01.03 | - | - | - | 6 | 6 | - | - | 2 |
| 1 | 4 | 72,307,616 | 247 | lncRNA01.04 | - | - | 3 | 6 | 6 | 5 | - | 2 |
| 1 | 4 | 72,307,630 | 14 | lncRNA01.05 | - | - | - | 6 | 6 | - | - | 2 |
| 1 | 5 | 77,440,144 | 5,132,514 | CSTA-5456-E03 | - | - | 4 | - | - | 6 | - | - |
| 1 | 5 | 77,440,155 | 11 | CSTA-5445-E03 | - | 5 | - | - | - | - | - | 3 |
| 1 | 5 | 77,440,159 | 4 | CSTA-5441-E03 | - | 5 | - | - | - | - | - | 3 |
| 1 | 5 | 77,442,145 | 1,986 | CSTA-3455-I01 | - | - | 4 | 7 | - | 6 | - | 3 |
| 1 | 5 | 77,442,831 | 686 | CSTA-2769-I01 | 2 | 6 | 0 | - | - | 6 | - | - |
| 1 | 5 | 77,444,900 | 2,069 | CSTA-0701-E01 | - | - | 4 | 7 | - | 6 | - | 3 |
| 1 | 5 | 77,444,931 | 31 | CSTA-0670-E01 | - | - | 4 | 7 | - | 6 | - | 3 |
| 1 | 5 | 77,567,379 | 122,448 | 6553B | 2 | 6 | 4 | - | - | 6 | - | 3 |
| 1 | 5 | 77,567,384 | 5 | 6553C | 2 | 6 | 0 | - | - | - | - | 3 |
| 1 | 5 | 77,862,924 | 295,540 | LAG3-0490-5P | 0 | 7 | 4 | 8 | - | 6 | - | 4 |
| 1 | 5 | 77,862,978 | 54 | LAG3-0544-5P | - | - | - | - | - | - | - | 3 |
| 1 | 5 | 77,863,256 | 278 | LAG3-0822-E02 | - | - | - | - | - | - | - | 3 |
| 1 | 5 | 77,863,325 | 69 | LAG3-0891-E02 | 3 | 7 | 4 | 8 | 7 | 6 | - | 0 |
| 1 | 5 | 77,863,508 | 183 | LAG3-1074-I02 | - | - | - | 0 | 8 | - | - | 3 |
| 1 | 5 | 77,864,317 | 809 | LAG3-1883-I04 | - | - | - | 0 | 0 | - | - | 0 |
| 1 | 5 | 77,864,660 | 343 | LAG3-2226-E05 | - | - | - | 0 | 7 | - | - | 3 |
| 1 | 5 | 77,864,774 | 114 | LAG3-2340-3P | 3 | 7 | 4 | 9 | 8 | 6 | - | 4 |
| 1 | 5 | 78,045,732 | 180,958 | C1S-00771-I01 | 0 | 8 | 5 | 10 | 9 | 7 | - | 5 |
| 1 | 5 | 78,045,886 | 154 | C1S-00925-I01 | - | 8 | - | 9 | 7 | 7 | - | 3 |
| 1 | 5 | 78,046,151 | 265 | C1S-01190-I02 | - | - | - | 9 | 8 | - | - | - |
| 1 | 5 | 78,046,415 | 264 | C1S-01454-I02 | - | 8 | - | 10 | 9 | 7 | - | 5 |
| 1 | 5 | 78,046,649 | 234 | C1S-01688-E03 | - | - | - | - | - | - | - | 4 |
| 1 | 5 | 78,047,963 | 1,314 | C1S-03002-E04 | - | 8 | 5 | 10 | 9 | 7 | - | 4 |
| 1 | 5 | 78,048,057 | 94 | C1S-03096-E04 | - | 8 | 5 | - | - | 7 | - | 3 |
| 1 | 5 | 78,048,417 | 360 | C1S-03456-E05 | - | 8 | 5 | 9 | 10 | 7 | - | 4 |
| 1 | 5 | 78,048,418 | 1 | C1S-03457-E05 | - | - | - | 9 | 10 | - | - | 0 |
| 1 | 5 | 78,049,446 | 1,028 | C1S-04486-E06 | - | - | - | - | - | - | - | 3 |
| 1 | 5 | 78,050,432 | 986 | C1S-05472-I08 | - | - | - | - | - | - | - | 3 |
| 1 | 5 | 78,050,684 | 252 | C1S-05724-I09 | - | - | - | 9 | 7 | - | - | - |
| 1 | 5 | 78,051,237 | 553 | C1S-06277-I10 | - | - | - | 9 | 7 | - | - | 3 |
| 1 | 5 | 78,051,418 | 181 | C1S-06458-E11 | - | - | - | 9 | 8 | - | - | - |
| 1 | 5 | 78,053,620 | 2,202 | C1S-08660-E12 | - | - | - | - | 0 | - | - | 3 |
| 1 | 5 | 78,053,783 | 163 | C1S-08823-E12 | - | - | - | 10 | 9 | - | - | 5 |
| 1 | 5 | 78,054,406 | 623 | C1S-09446-E12 | - | - | 5 | - | - | 7 | - | 3 |
| 1 | 5 | 78,055,130 | 724 | C1S-10170-3P | - | - | - | - | - | - | - | 4 |
| 1 | 7 | 104,379,743 | 26,324,613 | ADAMTS5-39503-E08 | - | 9 | 6 | 11 | 11 | 8 | - | 0 |
| 1 | 7 | 104,383,567 | 3,824 | ADAMTS5-35679-E07 | 4 | - | 7 | 11 | 12 | - | - | - |
| 1 | 7 | 104,383,654 | 87 | ADAMTS5-35592-E07 | - | - | - | - | - | 9 | - | 6 |
| 1 | 7 | 104,384,542 | 888 | ADAMTS5-34704-E06 | - | 9 | 7 | 0 | 12 | 9 | - | 6 |
| 1 | 7 | 104,385,136 | 594 | ADAMTS5-34110-E05 | - | 9 | 7 | 0 | 12 | 9 | - | 6 |
| 1 | 7 | 104,385,274 | 138 | ADAMTS5-33972-E05 | - | - | - | - | - | - | - | 6 |
| 1 | 7 | 104,386,766 | 1,492 | ADAMTS5-32480-I04 | 4 | 9 | - | 11 | 12 | - | - | 6 |
| 1 | 7 | 104,387,123 | 357 | ADAMTS5-32123-I03 | 4 | 9 | - | 11 | 12 | - | - | 6 |
| 1 | 7 | 104,392,241 | 5,118 | ADAMTS5-27003-E03 | - | - | - | 11 | 11 | - | - | - |
| 1 | 7 | 104,392,252 | 11 | ADAMTS5-26992-E03 | - | 9 | 6 | 0 | - | 8 | - | - |

| C | Q | bp | Distance | Marker | Line blocks | | | | | | | |
|---|---|---|---|---|---|---|---|---|---|---|---|---|
| | | | | | WL1 | WL2 | WL3 | WPR1 | WPR2 | WL4 | WL5 | RIR1 |
| 1 | 7 | 104,407,074 | 14,822 | ADAMTS5-12170-I02 | - | - | - | 11 | 11 | - | - | - |
| 1 | 7 | 104,407,555 | 481 | ADAMTS5-11689-I01 | - | - | - | 0 | 0 | - | - | - |
| 1 | 7 | 104,418,467 | 10,912 | ADAMTS5-00777-E01 | - | - | - | 11 | 11 | - | - | - |
| 1 | 7 | 106,098,899 | 1,680,432 | 1738A | - | 10 | 8 | 0 | 0 | 10 | - | 0 |
| 1 | 7 | 106,098,942 | 43 | 1738B | - | 10 | 8 | 0 | 0 | 10 | - | 0 |
| 1 | 9 | 173,657,608 | 67,558,666 | SUPT20H-00142-5P | - | - | - | 0 | 13 | - | - | 7 |
| 1 | 9 | 173,658,086 | 478 | SUPT20H-00620-E01 | - | - | - | 12 | 13 | - | - | 8 |
| 1 | 9 | 173,658,646 | 560 | SUPT20H-01179-I01 | 5 | 11 | 9 | 12 | 13 | 0 | 1 | 9 |
| 1 | 9 | 173,659,742 | 1,096 | SUPT20H-02275-E02 | - | - | - | - | - | - | - | 0 |
| 1 | 9 | 173,660,604 | 862 | SUPT20H-03138-I02 | 5 | 11 | 9 | - | - | 11 | 1 | 10 |
| 1 | 9 | 173,662,588 | 1,984 | SUPT20H-05122-I04 | 5 | 11 | 9 | - | - | 11 | 1 | 7 |
| 1 | 9 | 173,664,538 | 1,950 | SUPT20H-07072-I04 | - | - | - | 0 | 13 | - | - | 8 |
| 1 | 9 | 173,667,742 | 3,204 | SUPT20H-10276-E05 | - | - | - | 12 | 13 | - | - | 11 |
| 1 | 9 | 173,668,415 | 673 | SUPT20H-10949-E06 | - | - | - | 0 | 0 | - | - | - |
| 1 | 9 | 173,670,185 | 1,770 | SUPT20H-12719-E09 | 0 | 0 | - | 0 | 0 | 12 | - | 9 |
| 1 | 9 | 173,671,470 | 1,285 | SUPT20H-14004-I11 | - | - | - | 12 | 13 | - | - | 11 |
| 1 | 9 | 173,673,202 | 1,732 | SUPT20H-15736-E13 | - | - | - | 12 | 13 | - | - | 11 |
| 1 | 9 | 173,679,044 | 5,842 | SUPT20H-21579-I15 | - | - | - | 12 | 13 | - | - | 11 |
| 1 | 9 | 173,681,300 | 2,256 | SUPT20H-23835-I17 | - | 12 | - | 12 | 13 | 12 | - | 12 |
| 1 | 9 | 173,682,298 | 998 | SUPT20H-24833-I17 | - | - | - | - | - | - | - | 7 |
| 1 | 9 | 173,684,220 | 1,922 | SUPT20H-26875-I19 | - | - | - | 12 | 13 | - | - | 12 |
| 1 | 9 | 173,686,475 | 2,255 | SUPT20H-29130-I20 | - | - | - | 12 | 13 | - | - | 11 |
| 1 | 9 | 173,689,830 | 3,355 | SUPT20H-32485-I21 | - | - | - | 0 | 0 | - | - | 11 |
| 1 | 9 | 173,691,228 | 1,398 | SUPT20H-33883-E22 | 6 | 12 | - | 0 | 13 | 13 | - | 11 |
| 1 | 9 | 173,691,545 | 317 | SUPT20H-34200-E22 | 6 | 12 | - | - | - | 13 | - | 10 |
| 1 | 10 | 177,365,684 | 3,674,139 | FLT3_0386 | - | 13 | 10 | 0 | 14 | 14 | 2 | 13 |
| 1 | 10 | 177,365,756 | 72 | FLT3_0458 | - | 13 | 10 | - | - | - | 2 | - |
| 1 | 10 | 177,365,770 | 14 | FLT3_0472 | - | 13 | 10 | 0 | 14 | 14 | 2 | 0 |
| 1 | 10 | 177,366,081 | 311 | FLT3_0510 | - | 0 | - | 13 | 15 | 0 | - | 13 |
| 1 | 10 | 177,367,788 | 1,707 | FLT3_0553 | - | 13 | 10 | - | - | - | 2 | - |
| 1 | 10 | 177,369,603 | 1,815 | FLT3_0889 | - | 13 | 10 | 13 | 15 | 0 | 2 | 14 |
| 1 | 10 | 177,369,714 | 111 | FLT3_0922 | - | - | - | 13 | 15 | - | - | 0 |
| 1 | 10 | 177,374,264 | 4,550 | FLT3_1148 | - | - | - | - | - | - | - | 14 |
| 1 | 10 | 177,380,755 | 6,491 | FLT3_1380 | - | - | - | - | - | 0 | - | - |
| 1 | 10 | 177,385,869 | 5,114 | FLT3_2234 | - | 14 | 11 | 13 | 15 | 15 | - | 15 |
| 1 | 10 | 177,386,285 | 416 | FLT3_2266 | - | 14 | 11 | 13 | 15 | 13 | - | - |
| 1 | 10 | 177,386,295 | 10 | FLT3_2276 | - | 14 | 11 | - | - | 15 | - | 15 |
| 1 | 10 | 177,387,725 | 1,430 | FLT3_2663 | - | - | - | 13 | 15 | - | - | - |
| 1 | 10 | 177,388,614 | 889 | FLT3_2856 | - | 0 | 11 | - | - | 15 | - | - |
| 1 | 10 | 177,389,132 | 518 | FLT3_2962 | - | - | - | - | - | - | - | 14 |
| 1 | 11 | 196,386,357 | 18,997,225 | RELT-10463-E11 | - | 0 | - | 14 | - | - | - | 0 |
| 1 | 11 | 196,386,494 | 137 | RELT-10326-E11 | 7 | 15 | 0 | - | - | 16 | 3 | 0 |
| 1 | 11 | 196,386,900 | 406 | RELT-09920-I10 | - | 15 | 12 | 15 | 16 | 16 | - | 16 |
| 1 | 11 | 196,387,392 | 492 | RELT-09428-E09 | - | - | - | 14 | - | - | - | 16 |
| 1 | 11 | 196,389,295 | 1,903 | RELT-07525-E05 | - | - | - | 14 | - | - | - | 16 |
| 1 | 11 | 196,389,467 | 172 | RELT-07353-E04 | - | - | - | - | - | - | - | 0 |
| 1 | 11 | 196,389,592 | 125 | RELT-07228-E03 | - | - | - | 14 | - | - | - | 16 |
| 1 | 11 | 196,389,598 | 6 | RELT-07222-E03 | - | 15 | 12 | 15 | 16 | 0 | - | 0 |
| 1 | 11 | 196,390,599 | 1,001 | RELT-06221-I01 | 7 | 15 | - | - | - | 0 | 3 | - |
| 1 | 11 | 196,395,140 | 4,541 | RELT-01680-I01 | - | - | - | 15 | 0 | - | 3 | - |
| 2 | 12 | 633,591 | | 1749B | 0 | 0 | 0 | 0 | 0 | 0 | - | 1 |

| C | Q | bp | Distance | Marker | Line blocks | | | | | | | |
|---|---|---|---|---|---|---|---|---|---|---|---|---|
| | | | | | WL1 | WL2 | WL3 | WPR1 | WPR2 | WL4 | WL5 | RIR1 |
| 2 | 12 | 633,592 | 1 | 1749C | - | 0 | - | - | - | - | - | 1 |
| 2 | 12 | 788,532 | 154,940 | 6710 | - | - | 0 | 0 | 0 | 0 | - | - |
| 2 | 13 | 45,900,042 | 45,111,510 | TRANK1-9118-E21 | - | - | - | 1 | 1 | - | - | 2 |
| 2 | 13 | 45,900,106 | 64 | TRANK1-9054-E21 | 1 | - | 1 | 1 | 1 | - | 1 | 2 |
| 2 | 13 | 45,900,526 | 420 | TRANK1-8634-E21 | - | - | - | 1 | 1 | - | - | 2 |
| 2 | 13 | 45,900,538 | 12 | TRANK1-8621-E21 | - | - | - | 1 | 1 | - | - | 2 |
| 2 | 13 | 45,900,585 | 47 | TRANK1-8575-E21 | 0 | - | - | 1 | 1 | - | 1 | 2 |
| 2 | 13 | 45,900,688 | 103 | TRANK1-8472-E21 | - | - | - | 1 | 1 | - | - | 2 |
| 2 | 13 | 45,900,934 | 246 | TRANK1-8226-E21 | - | - | - | 1 | 1 | - | - | 2 |
| 2 | 13 | 45,901,252 | 318 | TRANK1-7908-E21 | 2 | 1 | 1 | - | - | - | - | - |
| 2 | 13 | 45,902,086 | 834 | TRANK1-7074-E21 | 2 | 1 | 1 | - | - | - | - | 3 |
| 2 | 13 | 45,902,476 | 390 | TRANK1-6684-E21 | 1 | 1 | 1 | - | - | - | 1 | 3 |
| 2 | 13 | 45,902,803 | 327 | TRANK1-6357-E21 | - | - | - | 1 | 1 | - | - | 2 |
| 2 | 13 | 45,906,374 | 3,571 | TRANK1-6094-E19 | - | 1 | - | - | - | - | - | - |
| 2 | 13 | 45,911,199 | 4,825 | TRANK1-5664-E15 | - | 2 | 2 | - | - | - | - | 3 |
| 2 | 13 | 45,911,265 | 66 | TRANK1-5598-E15 | - | - | - | 1 | 1 | - | - | 2 |
| 2 | 13 | 45,915,076 | 3,811 | TRANK1-5448-E14 | 1 | 1 | 1 | 1 | 1 | - | - | 2 |
| 2 | 13 | 45,915,618 | 542 | TRANK1-5353-E13 | - | 2 | 2 | 1 | 1 | - | - | 2 |
| 2 | 13 | 45,915,748 | 130 | TRANK1-5223-E13 | 1 | 1 | 1 | - | - | - | - | - |
| 2 | 13 | 45,916,424 | 676 | TRANK1-5163-E12 | - | - | - | 1 | 1 | - | - | 2 |
| 2 | 13 | 45,916,454 | 30 | TRANK1-5133-E12 | - | - | - | 1 | 1 | - | - | 2 |
| 2 | 13 | 45,917,333 | 879 | TRANK1-4254-E12 | - | - | - | 2 | 2 | - | - | - |
| 2 | 13 | 45,917,492 | 159 | TRANK1-4095-E12 | - | 2 | 2 | 2 | 2 | - | - | 2 |
| 2 | 13 | 45,917,699 | 207 | TRANK1-3888-E12 | - | 2 | 2 | 2 | 2 | - | - | 2 |
| 2 | 13 | 45,917,978 | 279 | TRANK1-3609-E12 | - | 2 | 2 | 2 | 2 | - | - | 4 |
| 2 | 13 | 45,918,480 | 502 | TRANK1-3107-E12 | - | - | - | 1 | 1 | - | - | 2 |
| 2 | 13 | 45,918,845 | 365 | TRANK1-2742-E12 | 1 | 1 | 1 | 2 | 2 | - | - | 3 |
| 2 | 13 | 45,921,445 | 2,600 | TRANK1-2196-E11 | 1 | 0 | 0 | 1 | 1 | - | - | 2 |
| 2 | 13 | 45,922,309 | 864 | TRANK1-2145-E10 | - | 2 | 2 | 2 | 2 | - | - | 3 |
| 2 | 13 | 45,925,740 | 3,431 | TRANK1-1689-E08 | - | 2 | 2 | 2 | 2 | - | - | 4 |
| 2 | 13 | 45,925,776 | 36 | TRANK1-1653-E08 | - | 2 | 2 | 2 | 2 | - | - | - |
| 2 | 13 | 45,925,812 | 36 | TRANK1-1617-E08 | - | 2 | 2 | 2 | 2 | - | - | - |
| 2 | 13 | 45,927,742 | 1,930 | TRANK1-1461-E07 | - | - | - | 2 | 2 | - | - | 4 |
| 2 | 13 | 45,932,906 | 5,164 | TRANK1-1011-E03 | - | 2 | 2 | 2 | 2 | - | - | 2 |
| 2 | 13 | 45,935,931 | 3,025 | TRANK1-0652-E01 | 1 | 2 | 2 | - | - | - | - | - |
| 2 | 14 | 107,692,346 | 61,756,415 | MD-07 | 0 | 0 | 0 | 0 | 0 | 0 | - | - |
| 4 | 18 | 8,508,640 | | 1462 | - | 0 | 0 | 0 | 0 | 0 | - | 0 |
| 4 | 19 | 87,451,251 | 78,942,611 | CTNNA2_085130_I10 | - | 1 | - | 0 | 1 | 0 | - | - |
| 4 | 19 | 87,455,417 | 4,166 | CTNNA2_080964_E09 | 1 | 2 | 1 | - | - | 1 | - | - |
| 4 | 19 | 87,461,520 | 6,103 | CTNNA2_074861_I07 | 1 | 2 | 1 | - | - | 1 | - | - |
| 4 | 19 | 87,469,358 | 7,838 | CTNNA2_067023_I06 | 1 | 0 | 1 | - | - | 1 | - | - |
| 4 | 19 | 87,479,459 | 10,101 | CTNNA2_056922_I05 | - | 3 | 2 | 1 | 2 | 0 | - | 0 |
| 4 | 19 | 87,482,774 | 3,315 | CTNNA2_053607_E04 | - | 3 | 2 | 1 | 2 | 0 | - | - |
| 4 | 19 | 87,483,115 | 341 | CTNNA2_053186_E03 | - | 1 | - | - | - | - | - | - |
| 4 | 19 | 87,518,550 | 35,435 | CTNNA2_017831_E02 | - | - | - | - | - | - | - | 0 |
| 4 | 19 | 87,535,920 | 17,370 | CTNNA2_000463_3P | 0 | 0 | 0 | 1 | 1 | 0 | - | - |
| 5 | 20 | 8,421,567 | | lncRNA06.01 | 1 | 1 | 0 | - | - | 0 | - | - |
| 5 | 20 | 8,422,079 | 512 | lncRNA06.02 | 1 | 1 | - | 0 | 0 | 0 | 0 | 1 |
| 5 | 20 | 8,544,549 | 122,470 | lncRNA07.03 | - | - | - | - | - | - | 1 | - |
| 5 | 20 | 8,544,933 | 384 | lncRNA07.04 | - | - | - | 1 | 1 | - | - | - |
| 5 | 20 | 8,545,143 | 210 | lncRNA07.05 | 0 | 2 | - | 1 | 1 | 0 | 1 | 1 |

| C | Q | bp | Distance | Marker | Line blocks | | | | | | | |
|---|---|---|---|---|---|---|---|---|---|---|---|---|
| | | | | | WL1 | WL2 | WL3 | WPR1 | WPR2 | WL4 | WL5 | RIR1 |
| 5 | 20 | 8,777,825 | 232,682 | lncRNA08.06 | 1 | 2 | 0 | 0 | 2 | - | 1 | 0 |
| 5 | 20 | 8,778,394 | 569 | lncRNA08.07 | - | - | - | - | 0 | - | - | 1 |
| 5 | 20 | 8,778,679 | 285 | lncRNA08.08 | - | - | - | 0 | 2 | - | - | - |
| 5 | 21 | 19,756,582 | 10,977,903 | lncRNA09.01 | - | 0 | - | 0 | - | 1 | 2 | - |
| 5 | 21 | 19,756,660 | 78 | lncRNA09.02 | 2 | - | 1 | 0 | 3 | 2 | - | 2 |
| 5 | 21 | 19,757,145 | 485 | lncRNA09.03 | - | - | - | 0 | 0 | 2 | - | 0 |
| 5 | 21 | 19,803,706 | 46,561 | lncRNA10.04 | 2 | 3 | 1 | 0 | 3 | 1 | - | 0 |
| 5 | 21 | 19,804,039 | 333 | lncRNA10.05 | - | 3 | - | 0 | 0 | 3 | - | 0 |
| 5 | 21 | 20,580,772 | 776,733 | lncRNA11.06 | - | 4 | - | 2 | 4 | 4 | 3 | 2 |
| 5 | 21 | 20,589,955 | 9,183 | lncRNA11.07 | - | 4 | 1 | 2 | 4 | 4 | - | 3 |
| 5 | 21 | 20,596,531 | 6,576 | lncRNA11.08 | - | 4 | - | 3 | 5 | 5 | 3 | 2 |
| 5 | 21 | 20,601,292 | 4,761 | lncRNA11.09 | - | 4 | 1 | 3 | 5 | 5 | - | 2 |
| 5 | 21 | 20,612,350 | 11,058 | lncRNA11.10 | - | - | - | 4 | 6 | - | - | - |
| 5 | 21 | 20,619,171 | 6,821 | lncRNA11.11 | - | 4 | - | 3 | 5 | 5 | - | 2 |
| 5 | 21 | 20,624,698 | 5,527 | lncRNA11.12 | - | 4 | 1 | 4 | 6 | 5 | - | 3 |
| 5 | 21 | 20,628,840 | 4,142 | lncRNA11.13 | - | 4 | 1 | 4 | 6 | 5 | - | 3 |
| 5 | 21 | 20,665,999 | 37,159 | lncRNA11.15 | - | 5 | - | 3 | 5 | 3 | 2 | 2 |
| 5 | 21 | 20,672,487 | 6,488 | lncRNA11.16 | - | 5 | - | 4 | 6 | 3 | 4 | 2 |
| 5 | 21 | 20,687,360 | 14,873 | lncRNA11.17 | - | 5 | - | 3 | 5 | 3 | 4 | 2 |
| 5 | 21 | 20,707,750 | 20,390 | lncRNA11.18 | - | 5 | 0 | 0 | - | 0 | 2 | 0 |
| 5 | 21 | 20,716,370 | 8,620 | lncRNA11.19 | - | 6 | 1 | 4 | 6 | 6 | 5 | 3 |
| 5 | 21 | 20,723,673 | 7,303 | lncRNA11.20 | - | 5 | - | 3 | 5 | 3 | 4 | 2 |
| 5 | 21 | 20,733,573 | 9,900 | lncRNA11.21 | - | 6 | 1 | 4 | 6 | 6 | 5 | 3 |
| 6 | 24 | 32,573,870 | | lncRNA17.14 | 1 | - | 1 | 1 | 1 | 1 | - | 0 |
| 6 | 24 | 32,579,493 | 5,623 | lncRNA17.15 | - | 1 | 2 | 1 | 1 | 1 | - | - |
| 6 | 24 | 32,583,416 | 3,923 | lncRNA17.16 | 1 | 1 | 2 | - | - | 2 | 1 | 0 |
| 6 | 24 | 32,588,194 | 4,778 | lncRNA17.17 | 1 | 1 | 1 | 1 | 1 | 2 | 1 | - |
| 6 | 24 | 32,590,409 | 2,215 | lncRNA17.18 | 1 | - | 1 | - | - | 1 | - | - |
| 6 | 24 | 32,634,574 | 44,165 | lncRNA12.01 | - | 2 | - | 2 | 2 | - | 1 | 1 |
| 6 | 24 | 32,635,144 | 570 | lncRNA12.02 | - | 3 | 2 | - | 2 | 2 | 1 | - |
| 6 | 24 | 32,635,255 | 111 | lncRNA12.03 | 1 | 3 | 3 | - | - | 3 | 1 | - |
| 6 | 24 | 32,639,215 | 3,960 | lncRNA13.04 | 1 | 0 | 3 | - | - | - | - | - |
| 6 | 24 | 32,642,143 | 2,928 | lncRNA13.05 | 1 | 3 | 3 | - | - | 3 | - | 2 |
| 6 | 24 | 32,644,318 | 2,175 | lncRNA13.06 | 1 | 0 | 2 | 2 | 0 | 3 | - | 1 |
| 6 | 24 | 32,698,298 | 53,980 | lncRNA14.07 | 1 | 4 | 2 | 1 | 1 | 3 | - | 2 |
| 6 | 24 | 32,698,617 | 319 | lncRNA14.08 | 1 | 4 | 2 | 1 | 1 | 3 | - | 2 |
| 6 | 24 | 32,712,118 | 13,501 | lncRNA15.09 | - | 4 | 1 | 1 | 1 | 4 | - | - |
| 6 | 24 | 32,713,410 | 1,292 | lncRNA15.10 | - | 2 | 1 | 1 | 1 | 4 | - | 3 |
| 6 | 24 | 32,803,271 | 89,861 | lncRNA16.11 | 1 | - | 2 | - | - | 3 | - | 3 |
| 6 | 24 | 32,803,462 | 191 | lncRNA16.12 | 1 | - | 2 | - | - | 3 | - | 3 |
| 6 | 24 | 32,803,638 | 176 | lncRNA16.13 | 1 | 0 | 1 | - | - | 3 | - | 3 |
| 7 | 25 | 13,676,411 | | 1659A | - | - | 0 | - | - | - | - | - |
| 7 | 25 | 13,676,441 | 30 | 1659B | 0 | - | 0 | 0 | 0 | - | 0 | 0 |
| 7 | 25 | 15,241,647 | 1,565,206 | MD-16 | - | - | - | 0 | 0 | - | 1 | - |
| 7 | 25 | 16,257,360 | 1,015,713 | 6624 | - | - | - | - | - | - | 1 | - |
| 7 | 25 | 16,772,561 | 515,201 | 1570 | 0 | - | - | 0 | - | - | 1 | 0 |
| 10 | 26 | 1,051,773 | | MD-20 | - | 0 | - | 0 | 0 | 0 | - | 0 |
| 10 | 26 | 2,033,349 | 981,576 | 6660 | - | 0 | - | 0 | 0 | 0 | - | 0 |
| 13 | 29 | 11,140,369 | | MD-22 | 0 | 0 | 0 | 0 | 0 | 0 | - | - |
| 13 | 29 | 12,164,061 | 1,023,692 | HAVCR1-0619-E01 | - | 0 | 1 | 1 | 1 | - | - | 1 |
| 13 | 29 | 12,164,859 | 798 | HAVCR1-1417-I01 | - | 1 | 2 | 2 | 2 | 1 | - | 1 |

| C | Q | bp | Distance | Marker | Line blocks | | | | | | | |
|---|---|---|---|---|---|---|---|---|---|---|---|---|
| | | | | | WL1 | WL2 | WL3 | WPR1 | WPR2 | WL4 | WL5 | RIR1 |
| 13 | 29 | 12,165,490 | 631 | HAVCR1-2049-E02 | - | - | 3 | - | - | - | - | - |
| 13 | 29 | 12,166,511 | 1,021 | HAVCR1-3069-I03 | - | 2 | 3 | 3 | 3 | 1 | 1 | - |
| 13 | 29 | 12,166,651 | 140 | HAVCR1-3209-I04 | - | 0 | 0 | - | - | 1 | - | - |
| 13 | 29 | 12,167,802 | 1,151 | HAVCR1-4360-I04 | - | 3 | 2 | 4 | 3 | - | - | 2 |
| 13 | 29 | 12,168,333 | 531 | HAVCR1-4891-I05 | - | - | - | 2 | 2 | 1 | 1 | 2 |
| 13 | 29 | 12,168,514 | 181 | HAVCR1-5072-E06 | - | 3 | 1 | - | - | 1 | - | - |
| 13 | 29 | 12,169,386 | 872 | HAVCR1-5944-E07 | - | - | - | - | - | 1 | 1 | 2 |
| 13 | 29 | 12,169,464 | 78 | HAVCR1-6022-I07 | - | 0 | 3 | - | - | - | - | - |
| 13 | 29 | 12,169,778 | 314 | HAVCR1-6336-I07 | - | 0 | 1 | 1 | 1 | - | - | 2 |
| 13 | 29 | 12,170,170 | 392 | HAVCR1-6728-I08 | - | 0 | 1 | 1 | 1 | 1 | - | 2 |
| 13 | 29 | 12,170,453 | 283 | HAVCR1-7011-E09 | - | 1 | 1 | 4 | 3 | - | 1 | 3 |
| 13 | 29 | 12,170,527 | 74 | HAVCR1-7085-E09 | - | 1 | 2 | 3 | 3 | 1 | 1 | - |
| 13 | 29 | 12,170,561 | 34 | HAVCR1-7119-E09 | - | 4 | - | - | - | - | - | - |
| 13 | 29 | 12,170,990 | 429 | HAVCR1-7548-E09 | - | - | - | 1 | 1 | - | - | 1 |
| 13 | 29 | 12,171,402 | 412 | HAVCR1-8072-E09 | - | 5 | - | 0 | 1 | - | 1 | 3 |
| 13 | 29 | 12,171,514 | 112 | HAVCR1-7960-E09 | - | 1 | 2 | 1 | 1 | - | 1 | - |
| 13 | 29 | 12,172,027 | 513 | HAVCR1-8585-E09 | - | 6 | 2 | - | - | - | - | - |
| 13 | 29 | 12,173,010 | 983 | HAVCR1-9558-E09 | - | 5 | - | 3 | 3 | - | 1 | - |
| 13 | 29 | 12,173,136 | 126 | HAVCR1-9694-3P | - | 2 | 3 | 3 | 3 | 1 | - | 1 |
| 13 | 29 | 12,174,321 | 1,185 | TIMD4-00530-5P | - | 5 | - | 3 | 3 | - | 1 | - |
| 13 | 29 | 12,174,479 | 158 | TIMD4-00688-I01 | - | 4 | 3 | - | - | - | - | - |
| 13 | 29 | 12,178,196 | 3,717 | TIMD4-04406-I01 | - | - | - | - | - | 1 | - | - |
| 13 | 29 | 12,182,052 | 3,856 | TIMD4-08262-I06 | - | 7 | 2 | 2 | 2 | - | 1 | 2 |
| 13 | 29 | 12,182,209 | 157 | TIMD4-08419-E07 | - | 7 | 2 | 2 | 2 | - | 1 | 2 |
| 13 | 29 | 12,182,218 | 9 | TIMD4-08428-E07 | - | - | - | - | - | 1 | - | - |
| 13 | 29 | 12,182,246 | 28 | TIMD4-08456-E07 | - | - | - | - | - | - | - | 2 |
| 13 | 29 | 12,182,885 | 639 | TIMD4-09095-I07 | - | 0 | - | 3 | 3 | 1 | 1 | 3 |
| 13 | 29 | 12,185,966 | 3,081 | TIMD4-12176-E11 | - | - | 3 | - | - | - | - | - |
| 13 | 29 | 12,186,115 | 149 | TIMD4-12325-I11 | - | 6 | 2 | 3 | 3 | - | - | 3 |
| 13 | 29 | 12,187,242 | 1,127 | TIMD4-13452-I13 | - | 6 | 2 | 3 | 3 | - | 1 | 1 |
| 13 | 29 | 12,188,510 | 1,268 | TIMD4-14720-E14 | - | 5 | 3 | 1 | 0 | 1 | 1 | 1 |
| 13 | 29 | 12,189,128 | 618 | TIMD4-15338-3P | - | 0 | - | 0 | - | - | - | - |
| 13 | 29 | 13,005,675 | 816,547 | MD-23 | 0 | 0 | 0 | 0 | 0 | - | - | 0 |
| 14 | 30 | 8,536,561 | | 1644A | - | - | - | - | 0 | - | 0 | - |
| 14 | 30 | 8,536,602 | 41 | 1644B | 0 | 0 | 0 | 0 | 0 | - | - | 0 |
| 14 | 30 | 9,146,787 | 610,185 | MD-28 | 0 | 0 | - | - | - | 0 | - | 0 |
| 14 | 30 | 9,343,521 | 196,734 | SOCS1-0315-5P | 0 | 0 | - | 0 | 0 | 0 | 1 | 0 |
| 14 | 30 | 9,343,654 | 133 | SOCS1-0448-5P | - | - | - | 1 | 1 | - | - | 1 |
| 14 | 30 | 9,344,120 | 466 | SOCS1-0914-E01 | 0 | 0 | - | - | - | 0 | 1 | - |
| 14 | 30 | 9,344,865 | 745 | SOCS1-1641-3P | - | - | - | 1 | 1 | - | - | 1 |
| 14 | 31 | 13,547,328 | 4,202,463 | 6643 | 0 | - | 0 | 0 | - | 0 | 0 | 0 |
| 14 | 31 | 14,468,249 | 920,921 | 1588 | - | - | - | 0 | 0 | - | - | 0 |
| 14 | 31 | 14,628,536 | 160,287 | 1715 | 0 | 0 | 0 | 0 | 0 | - | - | 0 |
| 16 | 32 | 2,357,051 | | MD-33 | - | - | - | - | 0 | 0 | - | - |
| 16 | 32 | 2,357,192 | 141 | MD-32 | - | 1 | - | 0 | 0 | 1 | - | 1 |
| 16 | 32 | 2,473,550 | 116,358 | IL4I1-0710-I01 | - | - | - | - | 1 | 1 | - | 1 |
| 16 | 32 | 2,475,052 | 1,502 | IL4I1-2212-E02 | - | 2 | - | - | 2 | 0 | - | 2 |
| 16 | 32 | 2,476,123 | 1,071 | IL4I1-3916-E04 | - | 3 | - | - | 2 | 0 | - | 2 |
| 16 | 32 | 2,476,132 | 9 | IL4I1-3925-E04 | - | 2 | - | - | 2 | 0 | - | 2 |
| 16 | 32 | 2,476,192 | 60 | IL4I1-3985-E04 | - | 1 | - | - | 0 | 0 | - | - |
| 16 | 32 | 2,476,467 | 275 | IL4I1-4260-E05 | 1 | 3 | - | - | 2 | 0 | - | 2 |

| C | Q | bp | Distance | Marker | Line blocks | | | | | | | |
|---|---|---|---|---|---|---|---|---|---|---|---|---|
| | | | | | WL1 | WL2 | WL3 | WPR1 | WPR2 | WL4 | WL5 | RIR1 |
| 16 | 32 | 2,476,563 | 96 | IL4I1-4356-I05 | 1 | 2 | - | - | 2 | 0 | - | 2 |
| 16 | 32 | 2,476,796 | 233 | IL4I1-4589-E06 | 1 | 2 | - | - | 2 | 0 | - | 2 |
| 16 | 32 | 2,476,811 | 15 | IL4I1-4604-E06 | 1 | 2 | - | - | 2 | 2 | - | 2 |
| 16 | 32 | 2,476,874 | 63 | IL4I1-4667-E06 | - | 2 | - | - | 0 | 2 | - | 3 |
| 16 | 32 | 2,477,464 | 590 | IL4I1-5256-E07 | 1 | 2 | - | - | 3 | 3 | - | 2 |
| 16 | 32 | 2,477,515 | 51 | IL4I1-5308-E07 | 1 | 2 | - | - | 3 | 3 | - | 2 |
| 16 | 32 | 2,477,572 | 57 | IL4I1-5365-E07 | 1 | - | - | - | 1 | 0 | - | 1 |
| 16 | 32 | 2,477,923 | 351 | IL4I1-5716-E07 | - | - | - | - | 0 | 0 | - | - |
| 16 | 32 | 2,550,271 | 72,348 | MD-30 | - | 1 | - | 1 | 0 | 4 | - | 1 |
| 16 | 32 | 2,551,587 | 1,316 | MD-29 | - | 1 | - | 1 | 4 | 0 | - | 1 |
| 16 | 32 | 2,597,942 | 46,355 | TAP1-3671-e07 | 1 | 2 | - | 1 | 1 | 4 | - | 2 |
| 16 | 32 | 2,597,967 | 25 | TAP1-3646-e07 | - | 3 | - | 1 | 5 | 4 | - | 2 |
| 16 | 32 | 2,598,372 | 405 | TAP1-3241-e06 | - | 1 | - | 1 | 0 | 0 | - | 2 |
| 16 | 32 | 2,599,123 | 751 | TAP1-2490-e05 | 1 | 3 | - | 1 | 5 | 0 | - | 2 |
| 16 | 32 | 2,599,748 | 625 | TAP1-1865-e04 | - | - | - | 0 | 0 | 0 | - | - |
| 16 | 32 | 2,600,004 | 256 | TAP1-1609-e03 | - | 2 | - | 1 | 4 | 0 | - | 2 |
| 16 | 32 | 2,600,170 | 166 | TAP1-1443-e02 | 1 | 2 | - | - | 6 | 0 | - | 3 |
| 16 | 32 | 2,600,920 | 750 | TAP1-0693-e01 | 1 | - | - | - | 6 | 0 | - | 3 |
| 17 | 33 | 3,938,971 | | TLR4-0562-5P | - | 1 | 1 | 1 | 1 | 1 | - | 0 |
| 17 | 33 | 3,940,192 | 1,221 | TLR4-1783-E02 | - | - | 1 | 1 | 1 | 1 | - | 1 |
| 17 | 33 | 3,941,832 | 1,640 | TLR4-3423-E03 | - | 2 | 2 | 1 | 1 | 2 | - | 1 |
| 17 | 33 | 3,941,957 | 125 | TLR4-3548-E03 | - | 2 | 2 | 1 | 1 | 2 | - | 1 |
| 17 | 33 | 3,942,076 | 119 | TLR4-3667-E03 | - | 2 | 3 | - | - | 2 | - | 2 |
| 17 | 33 | 3,942,761 | 685 | TLR4-4352-E03 | - | 2 | 2 | - | - | 2 | - | 2 |
| 17 | 33 | 3,943,548 | 787 | TLR4-5139-E03 | - | 2 | 3 | 2 | 2 | 2 | - | 2 |
| 17 | 33 | 3,944,254 | 706 | TLR4-5845-E03 | - | 2 | 3 | 2 | 2 | 2 | - | - |
| 17 | 33 | 3,944,711 | 457 | TLR4-6302-3P | - | 2 | 2 | 1 | 2 | 2 | - | - |
| 17 | 33 | 4,358,764 | 414,053 | BRINP1_1983 | - | 1 | 4 | 0 | 0 | 3 | - | 0 |
| 17 | 33 | 4,359,334 | 570 | BRINP1_1413 | - | - | - | 0 | 0 | - | - | 0 |
| 17 | 33 | 4,364,103 | 4,769 | BRINP1_1320 | - | - | - | 0 | 0 | 3 | - | - |
| 17 | 33 | 4,364,160 | 57 | BRINP1_1263 | - | 1 | 4 | 0 | 0 | 4 | - | 0 |
| 17 | 33 | 4,368,888 | 4,728 | BRINP1_1107 | - | 0 | 4 | - | - | 4 | - | - |
| 17 | 33 | 4,374,404 | 5,516 | BRINP1_0867 | - | - | - | 0 | 0 | - | - | 0 |
| 18 | 34 | 3,705,956 | | CD7_036 | - | - | - | - | - | - | - | 0 |
| 18 | 34 | 3,707,532 | 1,576 | CD7_276 | - | - | - | - | - | - | - | 0 |
| 24 | 35 | 4,813,725 | | 6612A | 1 | 1 | - | - | | 1 | - | - |
| 24 | 35 | 4,813,744 | 19 | 6612B | 1 | 1 | 0 | 0 | | 1 | 0 | - |
| 24 | 35 | 4,813,815 | 71 | 6612E | 1 | 1 | - | 0 | | 0 | - | - |
| 24 | 35 | 5,069,569 | 255,754 | 6671A | - | 0 | - | - | | - | - | 0 |
| 24 | 35 | 5,069,573 | 4 | 6671B | 0 | - | - | - | | - | - | - |
| 24 | 35 | 5,170,427 | 100,854 | 1745B | - | 0 | 0 | 0 | | - | - | 0 |
| 24 | 35 | 5,237,595 | 67,168 | TAGLN-2027-E02 | - | - | - | 0 | | - | - | - |
| 24 | 35 | 5,237,664 | 69 | TAGLN-2096-E02 | 2 | 2 | 1 | - | | 2 | 1 | - |
| 24 | 35 | 5,238,307 | 643 | TAGLN-2739-E05 | - | 2 | 0 | 1 | | 0 | - | 1 |
| 24 | 35 | 5,238,724 | 417 | TAGLN-3156-E05 | 2 | 0 | 1 | - | | - | 1 | 1 |
| 24 | 35 | 5,238,833 | 109 | TAGLN-3265-E05 | 2 | 0 | 0 | - | | 2 | - | - |
| 24 | 35 | 5,238,859 | 26 | TAGLN-3291-E05 | - | - | - | 1 | | - | - | 1 |
| 26 | 36 | 4,691,657 | | MD-44 | - | 1 | 1 | 1 | 1 | 1 | 1 | 1 |
| 26 | 36 | 4,755,292 | 63,635 | TREML2-6706-3P | - | 1 | 2 | 0 | 0 | 1 | - | 2 |
| 26 | 36 | 4,755,531 | 239 | TREML2-6467-E10 | - | 1 | 2 | 1 | 1 | 1 | - | 3 |
| 26 | 36 | 4,756,075 | 544 | TREML2-5923-E09 | - | - | 1 | - | - | - | 1 | - |

| C | Q | bp | Distance | Marker | Line blocks | | | | | | | |
|---|---|---|---|---|---|---|---|---|---|---|---|---|
| | | | | | WL1 | WL2 | WL3 | WPR1 | WPR2 | WL4 | WL5 | RIR1 |
| 26 | 36 | 4,756,562 | 487 | TREML2-5436-E08 | - | 1 | 3 | - | - | - | - | 2 |
| 26 | 36 | 4,756,606 | 44 | TREML2-5392-E08 | - | - | 1 | 1 | 1 | 1 | 1 | 4 |
| 26 | 36 | 4,756,618 | 12 | TREML2-5380-E08 | - | - | 1 | 1 | 1 | 1 | 1 | 4 |
| 26 | 36 | 4,756,951 | 333 | TREML2-5047-I07 | - | 1 | 1 | 0 | 0 | 1 | 1 | 0 |
| 26 | 36 | 4,757,206 | 255 | TREML2-4792-E07 | - | - | - | - | - | - | 1 | - |
| 26 | 36 | 4,759,369 | 2,163 | TREML2-2629-I04 | - | 1 | 3 | 1 | 1 | 1 | - | 1 |
| 26 | 36 | 4,759,648 | 279 | TREML2-2350-I04 | - | - | 1 | 1 | 1 | - | 2 | 0 |
| 26 | 36 | 4,759,889 | 241 | TREML2-2109-I03 | - | - | - | 0 | 0 | - | - | 1 |
| 26 | 36 | 4,759,960 | 71 | TREML2-2038-I03 | - | - | 1 | 1 | 1 | - | 1 | 3 |
| 26 | 36 | 4,761,468 | 1,508 | TREML2-0530-5P | - | - | 4 | 2 | - | 1 | 2 | 2 |
| 26 | 36 | 4,761,708 | 240 | TREML2-0290-5P | - | - | 4 | 2 | - | 1 | 2 | 2 |
| 27 | 37 | 3,923,054 | | 6573A | - | 1 | - | 1 | - | 1 | 1 | 1 |
| 27 | 37 | 3,923,070 | 16 | 6573B | - | 1 | 0 | 0 | 0 | 1 | 1 | 1 |
| 27 | 37 | 3,923,089 | 19 | 6573C | - | 1 | - | 1 | - | 1 | 1 | 1 |
| 27 | 37 | 4,040,797 | 117,708 | CD79B_1587 | 1 | 0 | - | - | - | 2 | - | 2 |
| 27 | 37 | 4,043,381 | 2,584 | CD79B_I2_110 | - | 2 | - | - | - | - | - | 2 |
| 27 | 37 | 4,045,076 | 1,695 | CD79B_0155 | - | 2 | - | - | - | - | - | 0 |
| 27 | 37 | 4,064,529 | 19,453 | MD-45 | 1 | - | 0 | - | - | 2 | 0 | 0 |
| 27 | 37 | 4,064,812 | 283 | MD-46 | - | - | - | 0 | 0 | 0 | - | 0 |

**Supplemental_Table_S12.** The number of all marker pairs tested in both lines (bottom left triangle), and among the tested pairs, pairs with $r^2 \geq 0.7$ shared by the two lines (upper right triangle). For example, Lines WL1 and WL2 shared 127 marker pairs, 40 of which had $r^2 \geq 0.7$.

| **Line** | **WL1** | **WL2** | **WL3** | **WPR1** | **WPR2** | **WL4** | **WL5** | **RIR1** |
|---|---|---|---|---|---|---|---|---|
| **WL1** | | 40 | 17 | 9 | 15 | 41 | 7 | 44 |
| **WL2** | 127 | | 36 | 29 | 34 | 66 | 12 | 70 |
| **WL3** | 73 | 264 | | 66 | 70 | 38 | 2 | 58 |
| **WPR1** | 31 | 358 | 295 | | 237 | 40 | 2 | 83 |
| **WPR2** | 85 | 494 | 356 | 2,069 | | 42 | 1 | 87 |
| **WL4** | 62 | 298 | 195 | 317 | 314 | | 7 | 82 |
| **WL5** | 17 | 49 | 6 | 25 | 13 | 54 | | 8 |
| **RIR1** | 118 | 610 | 296 | 1,122 | 1,291 | 529 | 60 | |

**Supplemental_Table_S13.** Genes and protein networks in $F_6$ QTLRs 4-5, ordered by location. Q - QTLR serial number (Supplemental_Table_S1); Start, End - gene coordinates in GRCg6a; Net - arbitrary number of a network seen in Figure 1, given by order of location of the first gene in the network (not by order of appearance in Figure 1): bolded - the network is comprised of genes from both QTLRs; B4-5 - location of the gene in the $F_6$ high LD blocks extending over the two QTLRs in the $F_6$ families.

| Q | Gene | Start | End | Net | B4-5 | Q | Gene | Start | End | Net | B4-5 |
|---|---|---|---|---|---|---|---|---|---|---|---|
| 4 | DUSP16 | 71,872,763 | 71,904,322 | 1 | + | 5 | ING4 | 77,734,990 | 77,749,769 | | + |
| 4 | CREBL2 | 71,960,834 | 71,970,851 | | + | 5 | ZNF384 | 77,755,770 | 77,781,779 | **4** | + |
| 4 | GPR19 | 71,980,272 | 71,988,795 | **2** | + | 5 | PIANP | 77,803,621 | 77,808,412 | | + |
| 4 | CDKN1B | 72,102,062 | 72,105,400 | | + | 5 | COPS7A | 77,817,350 | 77,820,482 | **4** | + |
| 4 | MRPS35 | 72,622,696 | 72,644,857 | **3** | + | 5 | MLF2 | 77,831,588 | 77,841,435 | **4** | + |
| 4 | MANSC4 | 72,649,269 | 72,664,039 | 1 | + | 5 | PTMS | 77,844,882 | 77,848,395 | | + |
| 4 | KLHL42 | 72,671,873 | 72,682,871 | **4** | + | 5 | LAG3 | 77,862,795 | 77,868,711 | **4** | + |
| 4 | PTHLH | 72,752,038 | 72,764,874 | **4** | + | 5 | CD4 | 77,873,930 | 77,885,897 | **4** | + |
| 4 | CCDC91 | 72,867,675 | 73,073,035 | **4** | + | 5 | GPR162 | 77,894,615 | 77,899,113 | **2** | + |
| 4 | FAR2 | 73,191,857 | 73,320,608 | | | 5 | P3H3 | 77,900,316 | 77,910,271 | | + |
| 5 | SLC2A14 | 75,548,453 | 75,558,943 | | | 5 | GNB3 | 77,917,296 | 77,922,102 | **4** | + |
| 5 | NANOG | 75,593,243 | 75,596,024 | | | 5 | CDCA3 | 77,922,769 | 77,924,912 | **4** | + |
| 5 | AICDA | 75,632,084 | 75,637,754 | **4** | | 5 | USP5 | 77,924,975 | 77,939,984 | **4** | + |
| 5 | MFAP5 | 75,647,640 | 75,660,701 | **4** | | 5 | TPI1 | 77,940,005 | 77,943,711 | **5** | + |
| 5 | RIMKLB | 75,676,381 | 75,723,185 | | | 5 | LRRC23 | 77,945,286 | 77,949,479 | | + |
| 5 | PHC1 | 75,865,514 | 75,884,654 | | | 5 | ENO2 | 77,952,924 | 77,962,832 | **5** | + |
| 5 | M6PR | 75,883,096 | 75,891,432 | **4** | | 5 | C1H12ORF57 | 77,991,895 | 77,993,033 | | + |
| 5 | OVST | 76,362,509 | 76,397,638 | **4** | | 5 | PTPN6 | 77,994,636 | 78,014,698 | **4** | + |
| 5 | MAN1A2 | 77,147,561 | 77,281,468 | | + | 5 | PHB2 | 78,015,335 | 78,020,461 | **6** | + |
| 5 | CD86 | 77,307,980 | 77,318,936 | **4** | + | 5 | EMG1 | 78,020,566 | 78,022,842 | **6** | + |
| 5 | CASR | 77,388,128 | 77,429,662 | **4** | + | 5 | LPCAT3 | 78,023,023 | 78,038,708 | **4** | + |
| 5 | CSTB | 77,433,979 | 77,438,548 | | + | 5 | C1S | 78,045,961 | 78,055,022 | 7 | |

| Q | Gene | Start | End | Net | B4-5 | Q | Gene | Start | End | Net | B4-5 |
|---|---|---|---|---|---|---|---|---|---|---|---|
| 5 | CSTA | 77,439,994 | 77,445,026 | | + | 5 | C1R | 78,058,485 | 78,065,761 | 7 | |
| 5 | CCDC58 | 77,454,326 | 77,464,115 | **3** | + | 5 | RBP5 | 78,070,071 | 78,071,702 | | |
| 5 | FAM162A | 77,463,747 | 77,472,850 | | + | 5 | CLSTN3 | 78,071,723 | 78,086,627 | | |
| 5 | KPNA1 | 77,477,877 | 77,514,773 | | + | 5 | PEX5 | 78,099,886 | 78,110,916 | | |
| 5 | FBXO40 | 77,526,063 | 77,541,005 | | + | 5 | EPHA1 | 78,162,527 | 78,195,976 | | |
| 5 | TAPBPL | 77,546,387 | 77,554,991 | | + | 5 | ZYX | 78,201,045 | 78,212,460 | | |
| 5 | gga-mir-6553 | 77,567,295 | 77,567,395 | | + | 5 | FAM131B | 78,257,533 | 78,259,472 | | |
| 5 | SCNN1A | 77,568,659 | 77,576,825 | | + | 5 | CLCN1 | 78,276,084 | 78,333,916 | | |
| 5 | VAMP1 | 77,579,970 | 77,584,568 | | + | 5 | CASP2 | 78,336,971 | 78,362,958 | | |
| 5 | MRPL51 | 77,585,845 | 77,587,396 | | + | 5 | TMEM139 | 78,364,767 | 78,369,487 | | |
| 5 | NCAPD2 | 77,587,505 | 77,611,310 | | + | 5 | RAP1GAP1 | 78,380,088 | 78,418,997 | | |
| 5 | SCARNA10 | 77,588,152 | 77,588,472 | | + | 5 | GSTK1 | 78,425,057 | 78,436,557 | | |
| 5 | CNP1 | 77,615,840 | 77,617,557 | | + | 5 | TAS2R40 | 78,481,425 | 78,482,360 | | |
| 5 | GAPDH | 77,619,214 | 77,623,350 | | + | 5 | TRPV6 | 78,604,630 | 78,633,785 | | |
| 5 | IFFO1 | 77,633,594 | 77,642,717 | | + | 5 | EPHB6 | 78,678,054 | 78,732,359 | | |
| 5 | NOP2 | 77,649,727 | 77,654,873 | | + | 5 | PRSS2 | 78,802,240 | 78,805,418 | | |
| 5 | LPAR5 | 77,694,435 | 77,702,350 | | + | 5 | PRSS3 | 78,879,681 | 78,923,402 | | |

**Supplemental_Fig_S1**. LD matrices found in the eight lines within and between the $F_6$ QTLRs, by line and chromosome. P - p-value of Trend association test (due to different QC criteria for the LD and the association analyses detailed in Methods, not all markers had p-values); B - high LD block ($r^2 \geq 0.7$) ordered by the location of the first marker, colored by block; Q - QTLR serial number (Supplemental_Table_S1), colored by QTLR; bp - the location on the GRCg6a genome assembly; Dist. - bp between the marker and the previous marker; No. - serial number of the marker; Marker/element, marker tested (Supplemental_Table_S11) colored by element: where the marker is within a candidate genomic element, the marker name includes the element name; LD values: red, $r^2 \geq 0.7$; pink, $0.15 \geq r^2 < 0.7$; white, $r^2 < 0.15$.

Chicken MD QTLRs
LD matrices by line.pc

#### LD blocks among QTLR elements in the eight lines

##### LD within one QTLR protein coding gene

Different high LD blocks were found even within QTLR genes over short distances. Appendix Table A presents an example of LD blocks within the gene *TRANK1* in QTLR 13 on chromosome 2 in Line WL1. Despite the short distances between markers within the same gene (390 to 14,486 bp), a complex pattern was found, with two LD blocks, one of which is fragmented around the other. There was high to complete LD between markers 5-8 and 13-36. These markers had practically no LD with markers 11 and 12, which had complete LD between them ($r^2 = 1.000$). Thus, in the gene *TRANK1* in Line WL1, Block 1 starts before, but ends after Block 2. The association test p-values [Smith et al., 2020] completely matched the LD blocks, with practically the same p-values in each block. This match was found in all other

QTLRs in all lines (e.g., Table 3, Appendix Tables 1 - 2). The p-values in Block 1 were all significant ($p \leq 0.05$), while Block 2 was not significant ($p = 0.500$). Thus, Markers 11 – 12 in Block 2, only 667 and 390 bp away from the nearest markers in Block 1, cannot be in linkage with the causative mutation(s) standing beyond the significance of Block 1.

**Table A.** LD within one QTLR protein coding gene. Line WL1, the gene TRANK1 in QTLR 13 on chromosome 2. P - p-values obtained by the association trend test [Smith et al., 2020], colored by the p-values; B - block serial number ordered by the location of the first marker, colored by the block; bp - marker location in GRCg6a; Dist. - distance between the present and previous marker in bp; No. - serial number of the marker; LD values: red, $r^2 \geq 0.70$; white, $r^2 < 0.15$.

| **P** | | | | | 3.0E-02 | 5.0E-02 | 5.0E-01 | 5.0E-01 | 3.0E-02 | 3.0E-02 | 3.0E-02 | 3.0E-02 | 3.0E-02 | 5.0E-02 |
|---|---|---|---|---|---|---|---|---|---|---|---|---|---|---|
| | **B** | | | | 1 | 1 | 2 | 2 | 1 | 1 | 1 | 1 | 1 | 1 |
| | | **bp** | | | 45,900,106 | 45,900,585 | 45,901,252 | 45,902,086 | 45,902,476 | 45,915,076 | 45,915,748 | 45,918,845 | 45,921,445 | 45,935,931 |
| | | | **Dist.** | | | 479 | 667 | 834 | 390 | 12,600 | 672 | 3,097 | 2,600 | 14,486 |
| | | | | **No.** | 5 | 8 | 11 | 12 | 13 | 18 | 20 | 28 | 29 | 36 |
| 3.0E-02 | 1 | 45,900,106 | | 5 | | | | | | | | | | |
| 5.0E-02 | 1 | 45,900,585 | 479 | 8 | 0.895 | | | | | | | | | |
| 5.0E-01 | 2 | 45,901,252 | 667 | 11 | 0.016 | 0.041 | | | | | | | | |
| 5.0E-01 | 2 | 45,902,086 | 834 | 12 | 0.016 | 0.041 | 1.000 | | | | | | | |
| 3.0E-02 | 1 | 45,902,476 | 390 | 13 | 1.000 | 0.895 | 0.016 | 0.016 | | | | | | |
| 3.0E-02 | 1 | 45,915,076 | 12,600 | 18 | 1.000 | 0.895 | 0.016 | 0.016 | 1.000 | | | | | |
| 3.0E-02 | 1 | 45,915,748 | 672 | 20 | 1.000 | 0.895 | 0.016 | 0.016 | 1.000 | 1.000 | | | | |
| 3.0E-02 | 1 | 45,918,845 | 3,097 | 28 | 1.000 | 0.895 | 0.016 | 0.016 | 1.000 | 1.000 | 1.000 | | | |
| 3.0E-02 | 1 | 45,921,445 | 2,600 | 29 | 1.000 | 0.895 | 0.016 | 0.016 | 1.000 | 1.000 | 1.000 | 1.000 | | |
| 5.0E-02 | 1 | 45,935,931 | 14,486 | 36 | 0.895 | 1.000 | 0.041 | 0.041 | 0.895 | 0.895 | 0.895 | 0.895 | 0.895 | |

**LD between two protein coding genes**

LD was also found between other types of QTLR elements. Table B presents such LD in Line WL2 between the QTLR protein coding genes *TLR4* and *BRINP1* in QTLR 33 on Chromosome 17. The first marker of *TLR4* has high LD to the first two markers of *BRINP1*, and the three markers are not linked to other markers of their own gene. The other six markers of *TLR4* form a complete LD block ($r^2$ = 1.000). Complexing it even further, the last marker of *BRINP1* (Marker 15) had low to moderate LD with all markers in QTLR 33, both genes included.

**Table B.** LD between QTLR protein coding genes. Line WL2, QTLR 33 on chromosome 17. P - p-values obtained by the association trend test [Smith et al., 2020], colored by the p-values; B - block serial number ordered by the location of the first marker, colored by the block; bp - marker location in GRCg6a; Dist. - distance between the present and previous marker in bp; No. - serial number of the marker, colored by QTLR protein coding genes [Smith et al., 2020]: green, TLR4; orange, BRINP1; LD values: Red, $r^2 \geq 0.7$; purple, $0.15 \leq r^2 < 0.70$; white, $r^2 < 0.15$.

| **P** | | | | | 2.9E-01 | 7.2E-01 | 7.2E-01 | 7.2E-01 | 7.2E-01 | 7.2E-01 | 7.2E-01 | 7.2E-01 | 1.6E-01 | 1.6E-01 | 8.4E-01 |
|---|---|---|---|---|---|---|---|---|---|---|---|---|---|---|---|
| | **B** | | | | 1 | 2 | 2 | 2 | 2 | 2 | 2 | 2 | 1 | 1 | |
| | | **bp** | | | 3,938,971 | 3,941,832 | 3,941,957 | 3,942,076 | 3,942,761 | 3,943,548 | 3,944,254 | 3,944,711 | 4,358,764 | 4,364,160 | 4,368,888 |
| | | | **Dist.** | | | 2,861 | 125 | 119 | 685 | 787 | 706 | 457 | 414,053 | 5,396 | 4,728 |
| | | | | **No.** | 2 | 4 | 5 | 6 | 7 | 8 | 9 | 10 | 11 | 14 | 15 |
| 2.9E-01 | 1 | 3,938,971 | | 2 | | | | | | | | | | | |
| 7.2E-01 | 2 | 3,941,832 | 2,861 | 4 | 0.135 | | | | | | | | | | |
| 7.2E-01 | 2 | 3,941,957 | 125 | 5 | 0.135 | 1.000 | | | | | | | | | |
| 7.2E-01 | 2 | 3,942,076 | 119 | 6 | 0.135 | 1.000 | 1.000 | | | | | | | | |
| 7.2E-01 | 2 | 3,942,761 | 685 | 7 | 0.135 | 1.000 | 1.000 | 1.000 | | | | | | | |
| 7.2E-01 | 2 | 3,943,548 | 787 | 8 | 0.135 | 1.000 | 1.000 | 1.000 | 1.000 | | | | | | |
| 7.2E-01 | 2 | 3,944,254 | 706 | 9 | 0.135 | 1.000 | 1.000 | 1.000 | 1.000 | 1.000 | | | | | |
| 7.2E-01 | 2 | 3,944,711 | 457 | 10 | 0.135 | 1.000 | 1.000 | 1.000 | 1.000 | 1.000 | 1.000 | | | | |
| 1.6E-01 | 1 | 4,358,764 | 414,053 | 11 | 0.830 | 0.102 | 0.102 | 0.102 | 0.102 | 0.102 | 0.102 | 0.102 | | | |
| 1.6E-01 | 1 | 4,364,160 | 5,396 | 14 | 0.830 | 0.102 | 0.102 | 0.102 | 0.102 | 0.102 | 0.102 | 0.102 | 1.000 | | |
| 8.4E-01 | | 4,368,888 | 4,728 | 15 | 0.157 | 0.661 | 0.661 | 0.661 | 0.661 | 0.661 | 0.661 | 0.661 | 0.171 | 0.171 | |
